# Cold adaptation in Upper Paleolithic hunter-gatherers of eastern Eurasia

**DOI:** 10.1101/2024.05.03.591810

**Authors:** Yusuke Watanabe, Yoshiki Wakiyama, Daisuke Waku, Guido Valverde, Akio Tanino, Yuka Nakamura, Tsubasa Suzuki, Kae Koganebuchi, Takashi Gakuhari, Takafumi Katsumura, Motoyuki Ogawa, Atsushi Toyoda, Soichiro Mizushima, Tomohito Nagaoka, Kazuaki Hirata, Minoru Yoneda, Takayuki Nishimura, Masami Izuho, Yasuhiro Yamada, Tadayuki Masuyama, Ryuzaburo Takahashi, Jun Ohashi, NCBN Controls WGS Consortium, Hiroki Oota

**Affiliations:** Department of Biological Sciences, Graduate School of Science, The University of Tokyo; Bunkyo-ku, Tokyo, Japan; Department of International Agricultural Development, Faculty of International Agriculture and Food Studies, Tokyo University of Agriculture; Setagaya-ku, Tokyo, Japan; Department of Anatomy, Kitasato University School of Medicine; Sagamihara, Kanagawa, Japan; Institute for Mummy Studies, Eurac Research; Bolzano, Italy; Department of Biosciences, Kitasato University School of Science; Sagamihara, Kanagawa, Japan; Center for the Study of Ancient Civilizations and Cultural Resources, College of Human and Social Sciences; Kanazawa University, Kanazawa, Japan; Advanced Genomics Center, National Institute of Genetics; Mishima, Shizuoka, Japan; Department of Anatomy, St. Marianna University School of Medicine; Kawasaki, Kanagawa, Japan; Aomori Public University; Aomori, Aomori, Japan; The University Museum, The University of Tokyo; Bunkyo-ku, Tokyo, Japan; Department of Human Life Design and Science, Faculty of Design, Kyushu University; Minami-Ku, Fukuoka, Japan; Faculty of Humanities and Social Sciences, Tokyo Metropolitan University; Hachioji, Tokyo, Japan; Educational Committee of Tahara City, Tahara, Aichi, Japan; Waseda University; Shinjuku-ku, Tokyo, Japan

## Abstract

Previous genomic studies understanding the dispersal of *Homo sapiens* have suggested that present-day East Eurasians and Native Americans can trace their ancestry to migrations from Southeast Asia. However, ineluctable adaptations during the Last Glacial Maximum (LGM) remain unclear. By analyzing 42 genomes of up to 30-fold coverage from prehistoric hunter-gatherers, Jomon, we reveal their descent from Upper Paleolithic (UP) foragers who migrated to and isolated in the Japanese archipelago during Late Pleistocene. We provide compelling evidence suggesting that these UP people underwent positive selection for cold environments, aiding their survival through the LGM facilitated by non-shivering thermogenesis and detecting it polygenically across multiple loci in the Jomon lineage. Our study pioneers the close estimation of the physiological adaptation of ancient humans by the paleogenomic approach.

## Introduction

The dispersal of *Homo sapiens* into Southeast Asia, estimated to have occurred at least 50 thousand years ago (ya) (*1*, *2*) following the out-of-Africa expansion, marked a significant peopling event in East Eurasian prehistory. Subsequently, divergent populations migrated northward, traversing eastern Eurasia (*3–5*). Recent studies leveraging ancient genome analysis of the Jomon people, prehistoric hunter-gatherers in the Japanese archipelago, have revealed intriguing insights into the ancestry of East Eurasian populations. Genetic affinity indicates a close relationship between a Jomon individual and Hoabinhian 8,000 year-old hunter-gatherers (*6*). Phylogenetic analysis shows that modern East Asians, Northeast Asians, and Native Americans diverged from Southeast Asians gradually, with a 40,000 year-old human skeleton discovered in Tianyuan Cave in China located at the root of East Asians, and the Jomon appearing as a very inner branch of the Tianyuan. Thus, the Jomon people can be modeled as a basal lineage of East Eurasians (*7*).

The Jomon genome is useful in unraveling the complex history of East Eurasian populations due to its geographical location. During the Late Pleistocene, present-day three main islands, Honshu, Shikoku, Kyushu (referred to collectively as “Hondo” in this paper; see Fig.S1) of the Japanese archipelago, was one island as called the Paleo-Honshu (P-Honshu) island, and almost bordered the Korean peninsula, with a very narrow strait. Hokkaido, now an island, was certainly connected to the East Eurasian continent at the mouth of Amur River, forming the Paleo-Sakhalin Hokkaido Kurile Peninsula (PSHK) (Fig. 1) (*8*, *9*). Cultural evidence indicates that the appearance of *Homo sapiens* equipped with small flake-based assemblage dates to ∼38,000 yr cal BP on P-Honshu Island (*10*, *11*). After the end of the LGM dating to 26,500-19,000 yr cal BP (*12*), rising temperatures and resulting sea level change led to fragmentation of the P-Honshu into three islands, and Hokkaido became isolated from the continent by the sea. This suggests the Jomon people, habitants in the Holocene, may represent an isolated group of ancient lineages originating from the East Eurasian continent in the Late Pleistocene and dating to the LGM when the Japanese archipelago separated from the continent. Examination of the Jomon genome may provide further insights into the genetic composition of the East Eurasian substratum population and the timing of these events.

**Fig. 1.**
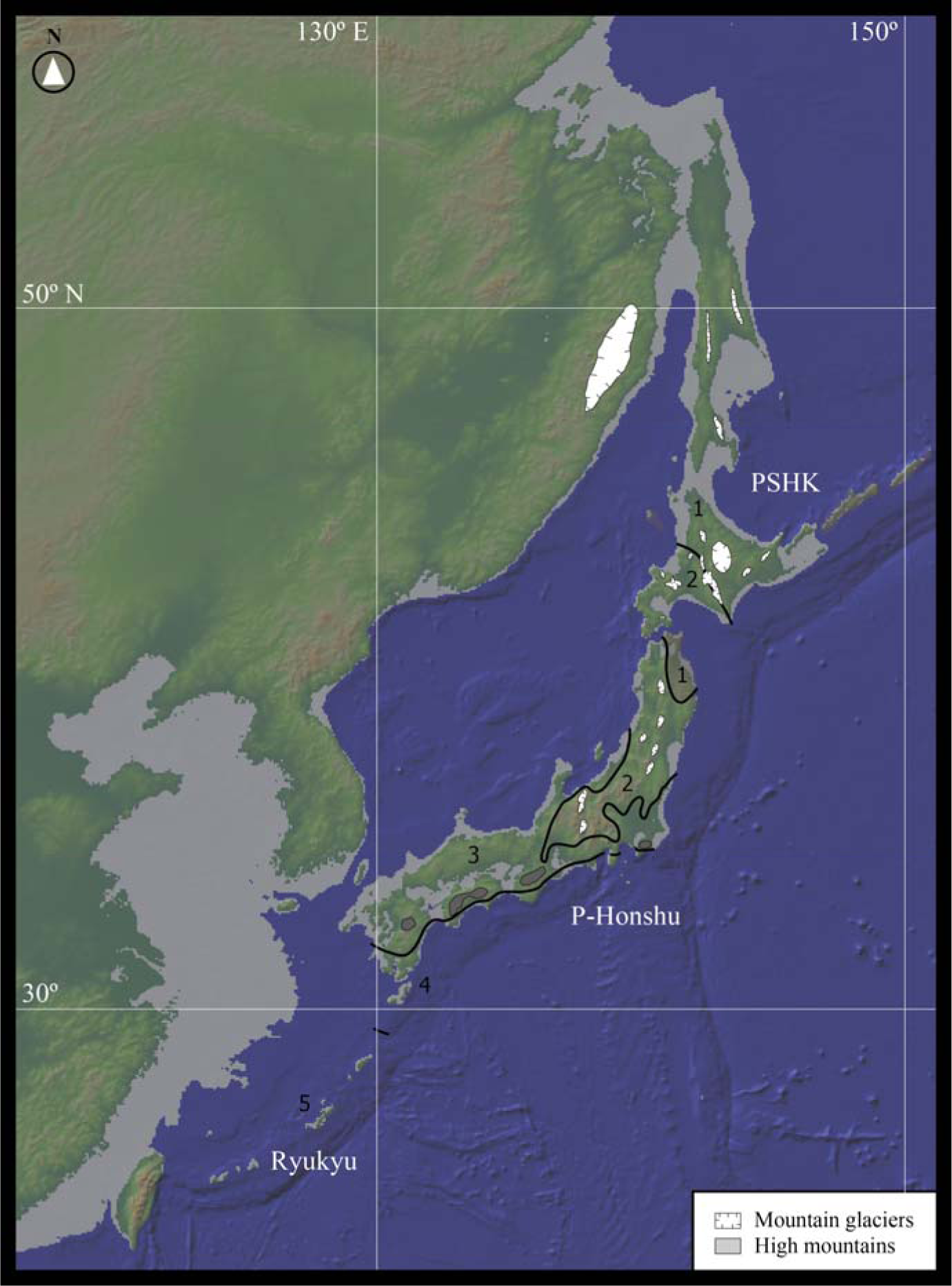
The map of the Japanese archipelago, showing the hypothesized coastline during the LGM (100 meters below the present sea level) and the distribution of vegetational areas during the LGM. 1: Open coniferous forest and grassland; 2: Cool-temperate coniferous forest; 3: Temperate pan-mixed forest; 4: Warm-temperate deciduous broadleaf and evergreen broadleaf forests; 5: Sub-tropical evergreen broadleaf forest and coniferous forest.

Although genomic evidence strongly suggests that modern Japanese ancestry results from hybridization between the Jomon and early farmers with paddy rice cultivation technology migrated from the East Eurasian continent about 3,000 ya (*7, 13–15*), it remains undetermined whether the Jomon people are direct descendants of Upper Paleolithic (UP) hunter-gatherers. Furthermore, if this proposed scenario is substantiated, a relatively rapid migration from Southeast Asia to East Asia would have occurred during the LGM although the mechanisms underlying this migration are unclear. Was it primarily a biological adaptation to the environment, or did cultural adaptations play a more significant role? Notably, the lack of genetic studies addressing this enigma underscores the need for further investigation. In this study, we present the results of our DNA extraction and sequencing of 25 Jomon bone samples, offering genomic insights and inferences into the peopling and adaptation of the UP East Eurasians. Considering these newly obtained genomes together with 17 previously reported Jomon samples, we conducted a comprehensive population genome analysis.

## Results and Discussion

### Genotype imputation of the Jomon people

We analyzed 42 Jomon genomes of newly sequenced (table S1 and S2) and previously reported (table S3) samples (*6*, *7*, *14–16*), including 3 high-coverage samples (two of which were newly sequenced), 18 samples with coverage ranging from 1x to 10x, and 21 low-coverage samples (< 1x). This study includes Jomon samples collected from various regions across the Japanese archipelago (fig. S2A). These ancient Jomon genomes were imputed and phased by GLIMPSE2 software (*17*), a tool designed for imputation of low-coverage genomic sequences. Given that approximately 20% of their genomic components of modern Hondo Japanese are reported to be derived from the Jomon people (*6*, *13*, *14*), and considering the absence of available direct descendants from the Jomon people in current populations today with the exception of Japanese archipelago populations, employing the modern Japanese as a reference was deemed necessary for the imputation of the Jomon genomes. For this purpose, we utilized a reference panel comprising the genomes of 9,290 modern Japanese individuals from the National Center Biobank Network (NCBN) (*18*), in addition to 2,482 individuals from the 1000 Genomes Project (1KG) (*19*) including 104 modern Japanese (JPT). We verified the imputation accuracy using high-coverage Jomon genomes [FUN23 (*14*) and IK2010-1] and found that the imputation accuracy, especially at polymorphic sites with minor allele frequency less than 1%, improved when employing a larger number of Japanese individuals in the reference panel (Fig. 2A).

**Fig. 2.**
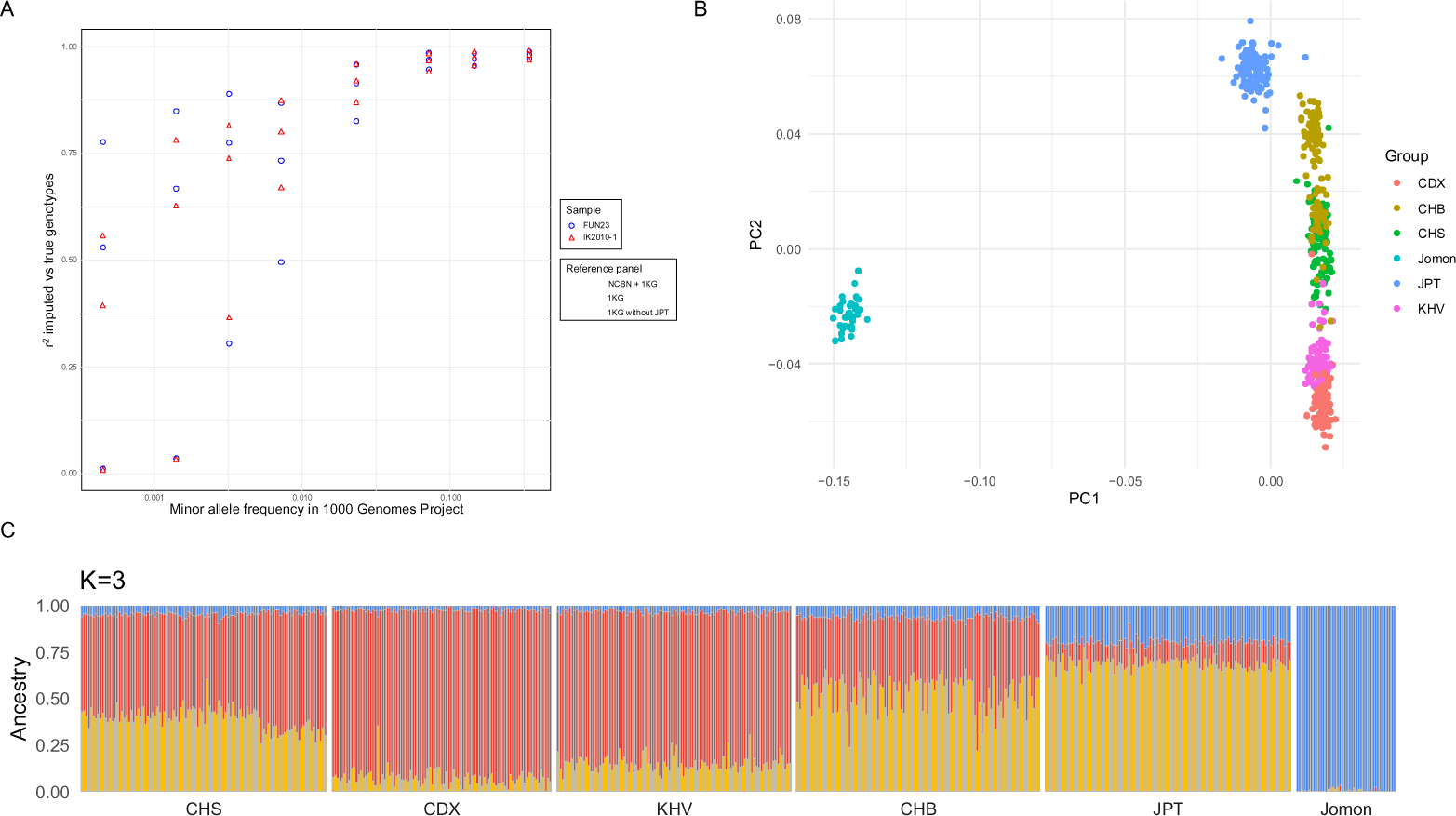
Imputation performance, principal component analysis (PCA) and ADMIXTURE analysis of the Jomon individuals. (A) Imputation accuracy (r^2^, y axis) on chromosome 22 for two high-coverage Jomon individuals (FUN23 (*14*) and IK2010-1) across the NCBN + 1KG, 1KG, 1KG without modern Japanese reference panels. Accuracy is plotted against log-scaled minor allele frequency in the 1KG panel (x axis). (B) PCA plot for 42 Jomon imputed genomes and 1KG East Eurasian genomes (modern Japanese, JPT; Han Chinese, CHB, Southern Han Chinese, CHS; Dai Chinese, CDX; Kinh Vietnamese, KHV) (C) ADMIXTURE analysis for 42 Jomon imputed genomes and 1KG East Eurasian genomes (K = 3)

Using the imputed genotypes of the Jomon people alongside contemporary East Asian genomes from the 1KG, we conducted principal component analysis (PCA) and ADMIXTURE analysis (*20*) (Fig. 2B and 2C). These analyses showed the 25 Jomon individuals in this study cluster with the 17 Jomon individuals previously published, further suggesting the Jomon people were geographically and genetically isolated from the modern East Asians in the continental region (*7*, *13–15*).

Subsequently, we investigated the lineage of the Jomon people through an analysis of 42 genomes. The earliest UP cultural evidence in PSHK Peninsula exhibits small-flake-based assemblages similar to those found on P-Honshu in terms of lithic technology by 30,000 yr cal BP (*21*). If UP hunter-gatherers independently migrated to PSHK and P-Honshu via two distinct routes—the mouth of Amur River and the Korean Peninsula, respectively—to form the Jomon people in each region, then we would expect distinct clusters of Jomon individuals on Hokkaido and Honshu on the phylogenetic tree. Conversely, a single cluster is more suggestive a unified entry route for the UP people transitioning into the Jomon people. Our phylogenetic analysis revealed that Jomon individuals on Hokkaido diverged from those in the northeastern region of Hondo (see fig. S2B). From this finding, it is plausible that a migration event occurred from Honshu to Hokkaido, casting doubt on the direct lineage of the Hokkaido Jomon (at least for the two individuals (*14*)) from the UP inhabitants of the area.

### Population history of the Jomon lineage since the UP

A previous study suggested that the Jomon people are direct descendants of basal East Eurasians, who are considered an ancestral population to modern East Asians, East Siberians, and Native Americans (*7*). This raises the question: What migration patterns and demographic history did the UP populations, ancestor(s) of the Jomon people, undergo subsequent to their divergence from the basal East Eurasians? Are the Jomon people direct descendants of UP people who had migrated northward from Southeast Asia to the Japanese archipelago and settled there some 38,000 years ago? To address these questions, we analyzed genomic data of 40 Jomon individuals having radiocarbon dating (the remaining two had no dates, see table S1 and S3). Employing the Relate software (*22*), we estimated population size changes within the Jomon lineage and modern East Eurasians.

We utilize multiple Jomon individuals and efficiently imputed single nucleotide polymorphisms (SNPs) with low minor allele frequencies, enabling a more comprehensive investigation into the post-divergence population history of the Jomon lineage compared to previous studies (*7*, *14*, *15*) (Fig. 3 AB, fig. S3, and S4). Fig. 3A depicts the overall fluctuation in population size of the Jomon lineage, encompassing diverse regional Jomon groups. Additionally, Fig. 3B illustrates the population size changes within each regional Jomon group. The light blue and red shadings in Fig. 3 correspond to the Last Glacial period (LGP) and the Jomon period, respectively. Figs. S3 and S4 depict the relative cross coalescence rate (RCCR), a metric calculated between pairs of populations, where a generation time at an RCCR of 0.5 indicates the divergence time between the two focal populations (*23*). Fig. S3 displays the RCCR between the Jomon lineage and modern East Asians from the 1KG, while Fig. S4 shows the RCCR between the Hondo and Hokkaido Jomon lineages. These analyses have yielded four significant findings: (1) The Jomon lineage diverged from continental East Asians approximately 27,000 to 19,000 years ago (fig. S3); (2) The population size of the Jomon lineage experienced a notable decline during the UP, persisting until the onset of the Jomon period (see Fig. 3A); (3) The population size of the Jomon lineage in Hondo, differing Hokkaido, began to recover at the beginning of the Jomon period (see Fig. 3B); (4) The Hokkaido Jomon lineage diverged from the Hondo Jomon lineage between 10,000 and 8,800 years ago, undergoing a distinct population history by a lack of subsequent recovery in population size during the Jomon period (Fig. 3B and fig. S4).

**Fig. 3.**
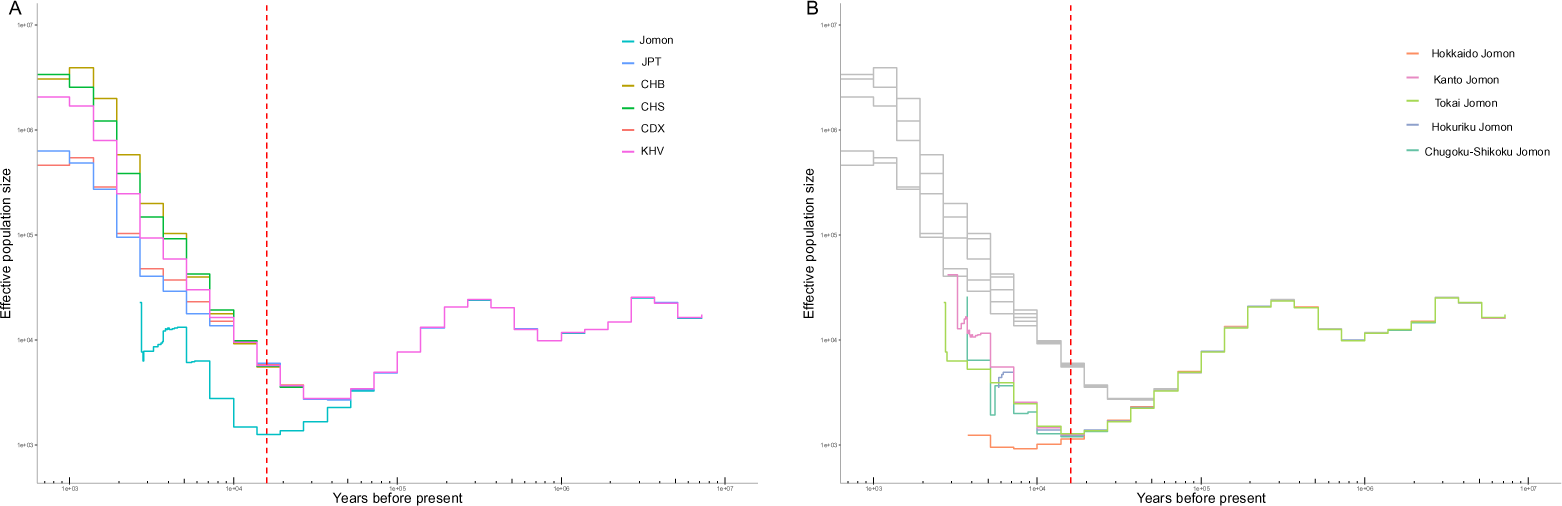
Effective population size change within the Jomon lineage and East Eurasians. We estimated population size using coalescence rate calculated by Relate-based genome-wide genealogies of 40 Jomon individuals who were subjected to radiocarbon dating and 1KG East Eurasians (JPT, CHB, CHS, CDX, KHV). (A) the overall population size change of the Jomon lineage. (B) the population size changes within each regional Jomon populations. The light blue shading: the Last Glacial period (10,000 YBP to 70,000 YBP); the red shadings: the Jomon period (3,000 YBP to 16,000 YBP).

The UP occupation began around 38,000 yr cal BP (*10*) and continued until about 15,000 yr cal BP on P-Honshu, extending from 15,000 to 12,000 yr cal BP on Hokkaido, south of PSHK (*21*, *24*). The UP period is divided into three sub-stages – early, middle, and late – characterized by lithic reduction patterns, toolkit compositions, stone-tool types, and date of the sites. Notably, the middle UP stage, spanning ∼30,000-20,000 yr cal BP, shows heightened mobility with blade-based backed point assemblages (*11*, *21*, *25*, *26*). Given the change observed in cultural evidences between the early and the middle UP (*10*, *21*, *25*), coupled with the estimated divergence time between the Jomon and continental East Asians (Fig. 3 and fig. S3), it is possible that the isolation of the Japanese archipelago from the East Eurasian continent just before or during the LGM resulted in a divergence between the UP populations on the Eurasian continent and those on the Japanese archipelago, leading to cultural transitions. The subsequent decline in the frequency of cultural occupation preceding to the Jomon period (*27*) closely aligns with our findings (Fig. 3), indicating a rapid decline in population size until the beginning of the Jomon period. Following an increase in effective population size after the LGM (Fig. 3), in the warming Japanese archipelago, the descendants of UP hunter-gatherers who had survived the ice age went to shape the Jomon culture, epitomized by its distinctive pottery (*28*). Moreover, the absence of recovery of the effective population size in the Hokkaido Jomon lineage is consistent with the line of cultural evidence (*9*, *25*, *29*, *30*), indicating a decrease in the intensity of cultural occupation in Hokkaido around 15,000 ya. Our findings are supported by archaeological evidence from the UP in the Japanese archipelago during this period (see Supplementary text). Furthermore, our findings may offer an avenue for testing hypotheses for the initial dispersal of the UP East Eurasians to the Americas (*29*, *31–33*) (see Supplementary text).

Subsequently, we conducted SPrime analysis (*34*) to identify archaic segments of Jomon and continental East Asians from the 1KG. We found no significant differences in the distribution of archaic segments between them, suggesting that hybridization events with Denisovans occurred prior to the divergence of the Jomon people from continental East Asian groups (see fig. S5 and Supplementary Text for detail). Thus, our analyses provide initial genetic evidence strongly supporting that the Jomon people are direct descendants of UP people who migrated into the Japanese archipelago and became isolated from the continent shortly prior to the end of the LGM and illustrates the value of Jomon genomes in understanding migration events and environmental adaptation.

### Cold adaptation in the Jomon lineage

To date, several previous studies have reported phenotypes derived from out-of-Africa populations that have been prevalent in East Eurasians (*35–44*). Table 1 showed the allele frequencies in the Jomon people that are strong association with phenotypes which are predominant in East Eurasians. We found that, aside from non-shivering thermogenesis (NST) (*43*, *44*), the Jomon people either lacked the alleles associated with phenotypes or possessed them at lower frequencies compared to modern East Asians from 1KG. Given that the Jomon people have been suggested to be direct descendants of basal East Eurasians, it is plausible that they did not frequently possess the derived phenotypes shared by subsequently diverged continental East Eurasians (e.g., thicker hair (*38*), shovel-shaped incisors (*39*), alcohol intolerance (*35*, *36*), light skin, iris color, freckles (*40–42*) and dry earwax (*37*)). Hence we were particularly interested in the GGTA haplotype, determined by four SNPs on the *UCP1* gene (rs3113195, rs12502572, rs1800592, rs4956451), leads to a thermogenic phenotype via NST (*43*, *44*), observed more frequently than in modern continental East Asians. A previous study reported a significant negative correlation between the frequency of the GGTA haplotype across geographic populations and the annual mean temperatures (*44*). By fitting the haplotype frequency of the Jomon people to the regression line as suggested by the previous study, we infer that the Jomon lineage was adapted to cold environments with an average temperature of about 5 °C (fig. S6).

**Table 1.** Allele frequencies in the Jomon people that are association with phenotypes predominant in East Eurasians.

| Phenotype | Gene | CHR | POS | rs ID | Ancestral | Derived | Allele Frequency associated with phenotype |  |  |  | Reference |
| --- | --- | --- | --- | --- | --- | --- | --- | --- | --- | --- | --- |
|  |  |  |  |  |  |  | Jomon | EAS | EUR | AFR |  |
| Thicker Hair<br>Shovel-shaped incisors | EDAR | 2 | 108897145 | rs3827760 | A | G | 0 | 0.87 | 0.01 | 0.003 | Fujimoto et al., (2008) |
|  |  |  |  |  |  |  |  |  |  |  | Kimura et al., (2009) |
| Alcohol intolerance | ADH1B | 4 | 99318162 | rs1229984 | C | T | 0 | 0.7 | 0.029 | 0.0015 | Edenberg and Borson (1997) |
|  | ALDH2 | 12 | 111803962 | rs671 | G | A | 0 | 0.17 | 0 | 0 | Harada et al. (1981) |
| Light skin color<br>Iris color | OCA2 | 15 | 27951891 | rs1800414 | T | C | 0.48 | 0.6 | 0 | 0.0008 | Yang et al. (2016) |
|  |  |  |  |  |  |  |  |  |  |  | Rawofi et al. (2017) |
| Dry earwax | ABCC11 | 16 | 48224287 | rs17822931 | C | T | 0.24 | 0.78 | 0.14 | 0.012 | Yoshiura et al. (2006) |
| Freckles | MC1R | 16 | 89919532 | rs2228479 | G | A | 0 | 0.29 | 0.069 | 0.0038 | Yamaguchi et al. (2012) |
| Light skin color |  | 16 | 89919746 | rs885479 | G | A | 0.35 | 0.61 | 0.07 | 0.0068 |  |
| Non-shivering thermogenesis<br>(NST) | UCP1 | 4 | 140554683 | rs3113195 | A | G | 0.67 | 0.47 | 0.67 | 0.24 | Hancock et al. (2011)<br>Nishimura et al. (2017) |
|  |  | 4 | 140563980 | rs12502572 | A | G | 0.67 | 0.48 | 0.67 | 0.24 |  |
|  |  | 4 | 140572807 | rs1800592 | T | C | 0.72 | 0.53 | 0.76 | 0.27 |  |
|  |  | 4 | 140580184 | rs4956451 | A | G | 0.72 | 0.53 | 0.67 | 0.1 |  |
| *The alleles denoted in bold in the Ancestral/Derived columns are associated with the East Asian characteristic phenotypes. |  |  |  |  |  |  |  |  |  |  |  |

Was the Jomon lineage adapted to cold environments? Under the assumption that the Jomon lineage had adapted to environments distinct from those of subsequently diverged East Eurasians, we performed a genome-wide selective sweep scan using cross-population nSL (XP-nSL) (*45*), with the Han Chinese from the 1KG as a proxy population. To investigate the phenotypes acquired through positive natural selection specific to the Jomon lineage, we devised a novel metric termed regional XP-nSL (rXP-nSL), integrating XP-nSL with modern Japanese GWAS data (*46–49*) (fig. S7). A positive rXP-nSL value means the positive natural selection for the Jomon lineage rather than continental East Eurasians. For each independent genetic region (i.e., without linkage disequilibrium) associated with traits, we calculated rXP-nSL, and regions displaying rXP-nSL greater than 6.566567, which is the 95-percentile value of 1,481 genetic regions in total among 62 trait GWASs, were deemed to exhibit significant signals of selective sweep in the Jomon lineage (fig. S8). To investigate the hypothesis of “cold adaptation in the Jomon lineage,” we focused on the rXP-nSL for genetic regions related to BMI and lipid metabolism (Fig. 4A). In assessing the overlapping effects on multiple traits within specific regions (including the *IRF2BP2* region in chromosome 1, the *APOB-TDRD15* region in chromosome 2, the *ZPR1-APOA5* region in chromosome 11, and the *ALDH1A2-LIPC* region in chromosome 15), we identified nine trait-associated regions across the whole genome that underwent positive natural selection, with implications for BMI or lipid metabolism in the Jomon lineage. Notably, our attention was drawn to the *FTO* gene region on chromosome 16 and the *ZPR1-APOA5* gene region on chromosome 11. The *FTO* gene region underwent positive natural selection for a haplotype associated with increased BMI (P = 4.9 × 10^-72^) (*46*), while the *ZPR1-APOA5* gene region exhibited selection for a haplotype linked to elevated triglyceride (TG) level (P < 1.0 × 10^-320^) (*48*).

**Fig. 4.**
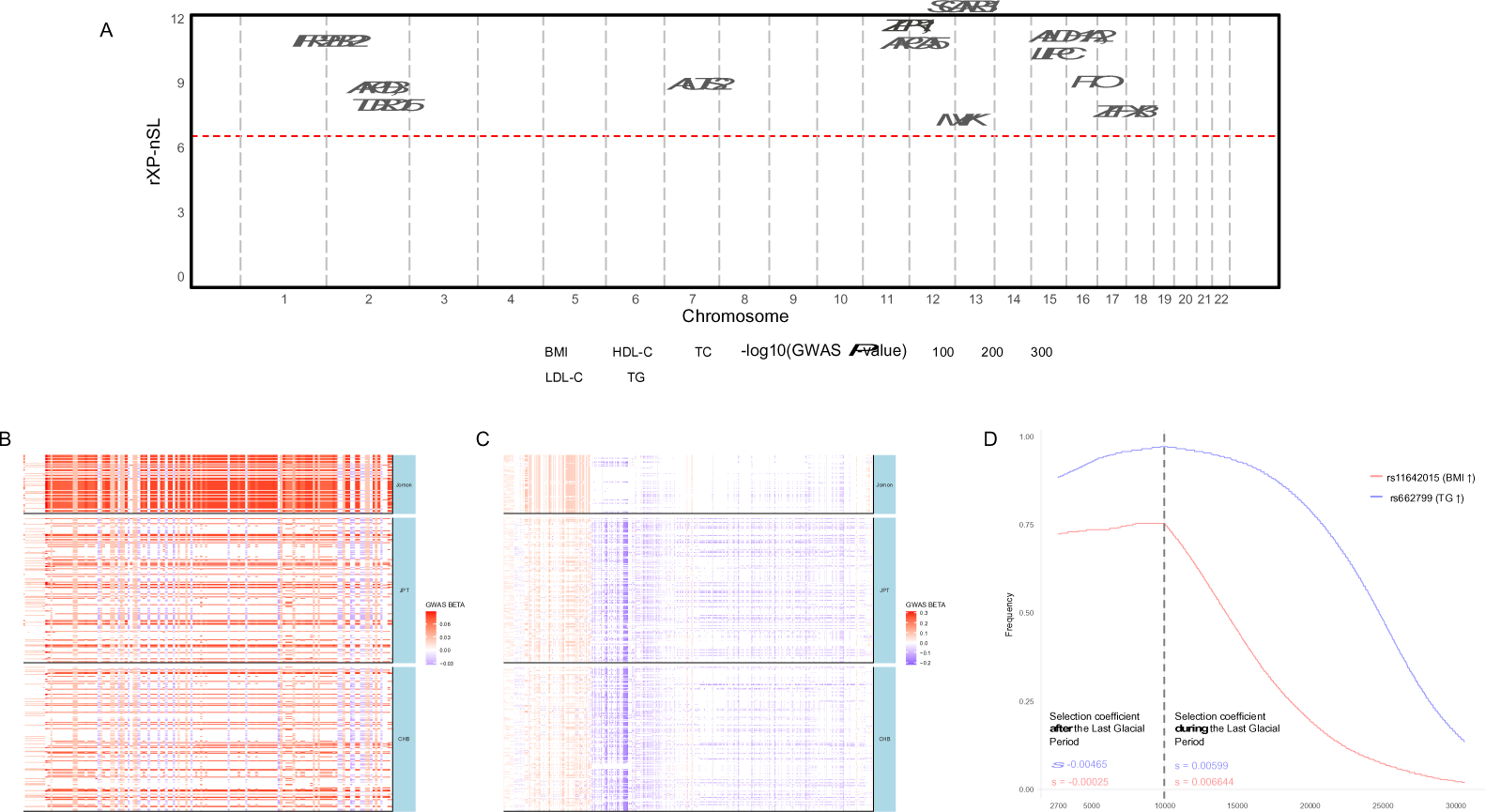
Selective sweep in genetic regions associated with cold adaptation. (A) rXP-nSL values for trait-associated regions which were detected by previous genome-wide association studies for BMI, HDL-C, LDL-C, total cholesterol (TC) and triglycerides (TG). rXP-nSL values are plotted against the physical positions on the chromosomes. We only illustrated regions with rXP-nSL > 0 where exhibit selective sweep signals for the Jomon people rather than continental East Eurasians. The 95th percentile of the empirical distribution are shown as red dashed horizontal line. Triangles mark regions with significant scaled XP-nSL values, and their orientation (upward or downward) signifies whether the selected haplotype in the Jomon people elevates or reduces the focal traits. (B) and (C) Haplotype structures of the *FTO* and *ZPR1-APOA5* gene regions among the Jomon people, CHB and JPT. Each column represents a trait-associated SNP site and each row represents a phased haplotype. The color of each cell represents the direction of effect for traits, and color intensity represents the β value for each effect allele identified by previous genome-wide association studies. (D) Allele frequency trajectories and selection coefficients (*s*) of the derived alleles for rs662799 and rs11642015. The derived allele for rs662799 in the *ZPR1-APOA5* region elevates TG, whereas the derived allele for rs11642015 elevates BMI. The dashed line shows the end of the Last Glacial period.

Figs. 4B and 4C illustrate the haplotype structures of the *FTO* and *ZPR1-APOA5* gene regions among the Jomon individuals, as well as CHB and JPT. In the *FTO* region, alleles highlighted in red (i.e., alleles associated with increased BMI) are more prevalent in the Jomon individuals (Fig. 4B) compared to CHB and JPT. Conversely, in the *ZPR1-APOA5* region, the Jomon individuals lack the accumulation of blue alleles (i.e., alleles associated with decreased TG) observed commonly in CHB and JPT, suggesting that the haplotypes prevalent in the Jomon individuals are associated with relatively higher TG levels compared to those in CHB and JPT (Fig. 4C). Among these genetic regions, two SNPs (rs662799 in the *ZPR1-APOA5* region and rs11642015 in the *FTO* region), which demonstrate the strongest association with their respective phenotypes, exhibited a higher frequency of the derived allele linked to increased BMI and TG levels in the Jomon people than in any other population worldwide from the 1KG (see Table 2). We further estimated the allele frequency trajectories and selection coefficients (*s*) of the derived alleles for rs662799 and rs11642015 using the CLUES2 software (*50*, *51*), utilizing genealogies obtained from Relate as inputs (Fig. 4D and fig. S9). The CLUES2 analyses unveiled robust evidence of positive natural selection for these alleles during the LGP (for rs662799, *s* = 0.0066 during the LGP and *s* = −0.00025 after the LGP; for rs11642015, *s* = 0.00599 during the LGP and *s* = −0.000465 after the LGP).

**Table 2.** Allele frequencies in rs662799 in the *ZPR1-APOA5* region and rs11642015 in the *FTO* region which exhibited the lowest P-values in the previous GWAS for their respective phenotypes (BMI and TG). For the Jomon people, we presented allele frequencies in 40 imputed genomes, as well as allele frequencies in 7 individuals with a depth greater than 5x, as genotype called by bcftools.

| Phenotype | Gene | CHR | POS | rs ID | Ancestral | Derived | Allele Frequency associated with phenotype |  |  |  |  | GWAS P-value |
| --- | --- | --- | --- | --- | --- | --- | --- | --- | --- | --- | --- | --- |
| | | | | | | | Jomon<br>(Imputed, n = 40) | Jomon<br>($\geq 5x$ depth,<br>called by bcftools, n = 7 ) | EAS | EUR | AFR | |
| Higher triglyceride level | *APOA5* | 11 | 116792991 | rs662799 | A | G | 0.89 | 0.79 | 0.29 | 0.08 | 0.12 | < 1.0E-320* |
| Higher BMI | *FTO* | 16 | 53768582 | rs11642015 | C | T | 0.73 | 0.57 | 0.17 | 0.43 | 0.06 | 4.9E-57† |
| *Kanai et al. (2018) |  |  |  |  |  |  |  |  |  |  |  |  |
| †Akiyama et al. (2017) |  |  |  |  |  |  |  |  |  |  |  |  |

Building upon our results, we consider into the cold adaptation of the UP hunter-gatherers of eastern Eurasia, the ancestors of the Jomon people through NST via *UCP1* on chromosome 4, *APOA5* on chromosome 11, and *FTO* on chromosome 16. NST is primarily produced by brown adipose tissue (BAT) in humans through the utilization of free fatty acids (FFA) derived from triglycerides (TG) (*52*). UCP1, highly and specifically expressed in the mitochondrial inner membrane of BAT, plays a crucial role in initiating NST (*52*). Notably, BAT levels increase during the winter season (*53*), and repeated exposure to cold also boosts BAT (*54*). Individuals with active BAT maintain a higher core body temperature with lower energy expenditure compared to those lacking BAT, who rely on shivering to cope with cold exposure (*55*), suggesting that NST from BAT might be a more efficient method of maintaining body temperature than shivering thermogenesis. A previous study has reported that accumulation of apoA5 protein, encoded by *APOA5*, within adipose cells accelerates intracellular TG metabolism and increases the expression level of the UCP1 protein (*56*). Therefore, if the Jomon lineage possessed genetic mechanisms that maintained higher blood TG levels through prevalent *APOA5* haplotypes among them (as illustrated in Fig. 4D), it is possible that an augmented supply of TG from the bloodstream to BAT facilitated heat generation by NST. A previous study found that the polygenic score (PS) for TG in the Jomon ancestry of modern Japanese was significantly higher than that of continental Asian ancestry and that regions with a higher proportion of Jomon ancestry among modern Japanese populations have higher obesity rates (*57*), lending further support to our findings. The *FTO* gene, is associated with obesity across various geographical populations in modern humans (*58*), harbors the rs1421085 T>C allele, whose functional role in NST has been corroborated by both in vivo and in vitro experiments. Introducing the rs1421085 T>C allele into mice has been shown to enhance thermogenesis via upregulating *Fto* expression and promoting the maintenance of body temperature under cold exposure (*59*), indicating that the rs1421085 C allele is a functional variant promoting brown-fat thermogenesis. Interestingly, the rs1421085 C allele was encompassed within the haplotype subject to positive natural selection in the Jomon people (Fig. 4B), with allele frequencies in the Jomon people and worldwide populations are as follows: 0.725 in the Jomon people, 0.168 in East Asians (EAS), 0.43 in Europeans (EUR), and 0.056 in Africans (AFR). Thus, it is plausible that the higher allele frequency of rs1421085 C in the Jomon people likely facilitated NST through increased expression of *FTO*. In light of these findings, coupled with the timing of natural selection (Fig. 4D), we propose that the UP hunter-gatherers of eastern Eurasia were cold adapted through brown-fat thermogenesis involving *UCP1*, along with *APOA5* and *FTO* haplotypes, during the LGP and before the onset of the Jomon period.

Human cold adaptation is classified into several patterns among local populations (*60*). These adaptation patterns are primarily categorized as metabolic adaptation, insulative adaptation, and hypothermic adaptation. Notable examples include the Inuit peoples exhibiting metabolic adaptation by enhancing thermogenesis through high-calorie food intake to maintain body temperature in extreme cold environments (*61*); indigenous peoples of northern coastal Australia demonstrating insulative adaptation with unchanged thermogenesis and lower skin temperature to preserve core body temperature in cold conditions (*62*); and the Bushmen of the Kalahari Desert manifesting hypothermic adaptation by lowering both core body and skin temperatures through reduced thermogenesis (*63*). These diverse adaptation patterns are influenced by factors including cold stress intensity, hormonal fluctuations, physical attributes, and dietary practices that have shaped and continue to shape our evolutionary history. Although several polymorphisms have been proposed as candidates for carriers of cold adaptation through NST (*59*, *64*), it may be that the Jomon lineage represents the first instance where signals of cold adaptation through NST have been polygenically detected, observed consistently across multiple loci.

Additional evidence supporting cold adaptation through NST in the Jomon lineage is the similarity of allele frequencies to those observed in the Greenland Inuit. Notably, the allele frequencies observed in the Jomon people for SNPs located in *FADS1, FADS2, FADS3*, genes known to have undergone positive natural selection in the Greenland Inuit (*65*), were found to be significantly closer to those of Greenlandic Inuit than to Han Chinse, who share a more recent common ancestor with the Jomon lineage (fig. S10, table S4). Through dietary change, the Jomon lineage may have adapted to cold environments similar to Greenlandic Inuit cold adaptation to diets rich in omega-3 polyunsaturated fatty acids (PUFAs) abundant in seafood (*65*). A previous study demonstrated that modern hunter-gatherers in cold climate regions derive approximately 85% of their nutrition from meat through hunting or fishing (*66*). Another study has reported traces of both marine and freshwater resources on Jomon pottery, dating from about 15,000 to 11,800 years ago, found across the Japanese archipelago (*67*). Craig et al. (2013) concluded that the Incipient Jomon people, foragers at the end of the glacial period, primarily depended on marine and freshwater diets. Given the diets of later Jomon populations were not solely dependent on marine products (*68–70*), it is likely that dietary adaptation of the Jomon lineage originated not during the Jomon period but among the UP hunter-gatherers of eastern Eurasia during the LGM. This dependence on marine diets may have persisted into the Incipient Jomon period. Adaptation to cold environments through NST offers distinct advantages under severe cold conditions, particularly when dietary lipids and proteins are readily available from meat and/or seafood. This is primarily because thermogenesis is the only generative mechanism to raise core body temperature. By contrast, insulative or hypothermic adaptations conserve energy metabolism, and likely only help to maintain body temperature through heat loss prevention. Given the high prevalence of thermogenic haplotypes and haplotypes adapted to a diet rich in PUFAs among the Jomon people, it is hypothesized that their UP hunter-gatherer ancestors had access to abundant high-fat, high-protein marine resources, akin to those utilized by modern Inuit. In light of these conditions, metabolic adaptation represents a more likely explanation compared to insulative or hypothermic adaptations.

A paleogenomic study has indicated that modern East Asians are descendants from people who migrated south of the Himalayas to reach the eastern side of the Eurasian continent (*7*). It is proposed that the common ancestors of East Eurasians, including the Jomon lineage, initially settled in warmer rather than colder environments, likely around Southeast Asia. Given the early divergence and reduction in population size during the LGM as depicted in Fig. 3, we hypothesize that the UP hunter-gatherers, as direct ancestors of the Jomon people, migrated northward across the East Eurasian continent and reaching the Japanese archipelago just before or during the LGM. Considering the higher selection coefficients observed in the *FTO* and *APOA5* regions during the LGP compared to those after the LGP in the Jomon lineage as shown in Fig. 4D, it suggests that the northward migration of the ancestral UP hunter-gatherers, who in part become the Jomon people, necessitated adaptation to colder environments compared to diverged continental East Eurasian populations.

In summary, our analysis of 42 Jomon genomes argues in support of their descent from populations diverging from the continental UP, settling in the Japanese archipelago ∼27-19,000 ya. Our findings affirm that the Jomon, as an isolated group, preserved a genome rich in prototypes of the UP for over 10,000 years, facilitated by the geographic isolation of the archipelago from the continent following the sea-level rise at the end of the Last Glacial Period (around 16,000 ya). These results highlight the significance of the Jomon genome as a valuable repository of UP traits from East Eurasia. Through our analysis of the Jomon genome, we present multiple lines of evidence for cold adaptation during the UP of East Eurasia. Given the stark differences in dietary habits and living environments between ancient times and today, our understanding of ancient human physiological phenotypes remains largely speculative. Regarding the Japanese archipelago, since almost no organic materials including human bones during the UP periods have remained today, researchers know absolutely nothing about the diet of the UP inhabitants. This study pioneers a paleogenomic approach to closely estimating the physiological functions, dietary habits and adaptations of the UP humans, drawing on comprehensive genetic evidence and corresponding phenotypes. We anticipate further advancements in this field.

## Materials and Methods

### Archaeological Samples

This study used 25 human remains from five sites assigned to the Middle, the Last, and the Final Jomon periods [the Middle: 5,400–4,400 yr cal BP; the Late: 4,400–3,300 yr cal BP; the Final: 3,300-2,300 yr cal BP (*71*)]. Below are the comprehensive archaeological details of the aforementioned samples.

#### Shell-mounds in Ichihara City

We examined the human remains from three shell-mounds, Kikumatenaga, Gionbara, and Saihiro, located in Ichihara City, Chiba Prefecture, overlooking Tokyo Bay. Excavations at the Kikumatenaga (KT) Shell-mound were conducted between 1972 and 1995, unearthing approximately 80 Jomon individuals. Radiocarbon dating placed these individuals in 5,280–3,490 yr cal BP (table S1) (*72*). Similarly, the Gionbara (GB) Shell-mound underwent five surveys between 1977 and 1995, yielding 112 human skeletal remains from the shell layer of the Late Jomon period. Archaeological analysis suggests that both sites were utilized from the Late Jomon period to the Final Jomon period, with individuals dated to between 4,295–3,826 yr cal BP (table S1) (*72*). Additionally, the Saihiro (SH) Shell-mound underwent seven surveys between 1972 and 1987 as part of road construction, unearthing 72 human skeletons. Archaeological analysis of these remains indicated usage of this site from the end of the Middle Jomon period to the early half of the Final Jomon period, with individuals dated to between 4,404–3,779 yr cal BP (table S1) (*72*).

#### Kosaku Shell-mound in Funabashi City

Situated in Funabashi City, Chiba Prefecture, overlooking Tokyo Bay, the Kosaku (KS) Shell-mound is has been under excavation since 1926, resulting in the collection of approximately 100 human skeletons from this site. Among them, the subject of this study, KS007, is an infant discovered cradled in the chest of a woman in her prime. Based on pottery chronology, this site is estimated to the Late Jomon period. Radiocarbon dating of KS007 places its age at 4,420 (95.4%) 4,291 yr cal BP, consistent with the archaeological contexts of this layer.

#### Ikawazu Shell-mound in Tahara City

Situated in Tahara City, Aichi Prefecture, near the center of the Japanese main island (Honshu), the Ikawazu (IK) Shell-mound has been a focal point of excavation since its initial exploration in 1918, yielding over 100 human bones accompanied by Jomon pottery. Based on pottery chronology, this site is classified as the Late–Final Jomon period. The individuals examined in this study were found in association with the type of Gokan-no-mori pottery, characterized by the absence of evidence for rice cultivation. Our analysis confirmed their affiliation with Jomon period, marked by typical Jomon cultural traits. Radiocarbon dating of five individuals showed a range of 2,848–2,712 yr cal BP (table S1), with the four of these individuals previously dated by Yamada et al., (2022) (*73*).

### Gelatin extraction and C14 dating

Collagen was extracted from each sample and purified gelatin was obtained using protocols from previous studies (*74*, *75*). Radiocarbon dating was carried out using a compact Accelerator Mass Spectrometry (AMS) at The University Museum, The University of Tokyo. Calibrated radiocarbon ages were calculated by referencing the IntCal20 and Marine 20 datasets (*76*, *77*), using OxCal 4.4 software (*78*).

Given the shell-mound origin of our samples, predominantly sourced from marine resources, stable isotope ratios of carbon and nitrogen were measured from extracted gelatins. This allowed for correction of the marine reservoir effect, accounting for variations in the intake ratio of marine fish and shells. A regional correction value (ΔR) specific to Tokyo Bay (−98 ± 37 years) was applied to Marine20, based on analysis of a shell specimen collected in 1882 AD (534 ± 36 BP) (*79*).

For Ikawazu Jomon, the contribution of amino acids from marine resources was estimated by considering the spectrum between terrestrial herbivores and marine fish. This analysis suggested that approximately 50% of the diet at Ikawazu Jomon was derived from marine sources.

### DNA extraction

DNA extraction and library preparation were performed in a dedicated clean room exclusively designed for ancient DNA analyses, installed in the Department of Anatomy, Kitasato University School of Medicine, or the Department of Biological Sciences, Graduate School of Science, University of Tokyo. DNA extraction followed protocols established in the previous studies (*7*, *80–82*). Petrous bones were cut using a UV-irradiated disc and drill to access the inner part. Bone pieces were treated with 2 mL of lysis buffer containing 20 mM Tris HCl (pH 7.5), 0.7% N-lauroylsarcosine, 0.5 M EDTA (pH 8), and 0.65 U/mL recombinant Proteinase K, shaken at 900 rpm and 50 °C in a Thermomixer (Eppendorf) for 15 minutes. After discarding the supernatant, 2 mL of fresh lysis buffer was added, and the tubes were incubated for > 16 h with shaking at 900 rpm and 50 °C in a Thermomixer. Subsequently, the samples were centrifuged at 13,000 x g for 10 min, and the supernatants were transferred to ultrafiltration tubes (Amicon® Ultra-4 Centrifugal Filter Unit 10K, Merck), diluted with 2 ml TE (pH 8.0) and centrifuged at 2,300 x g until final concentrations of ∼100 µL were obtained. These concentrations were then transferred to silica columns (MiniElute PCR Purification Kit, QIAGEN) and purified according to the manufacturer’s instructions, except for elution with EBT buffer (EB buffer with 0.005% TWEEN 20). The EBT buffer was preheated to 60°C and added 60 µL. An extraction blank was incubated at this step to check for cross-and background contamination and processed in parallel with the sample. DNA extracts were qualified by Qubit 4.0 (Thermo Fisher Scientific) and Bioanalyzer (Agilent).

### Library construction and sequencing

Library construction was carried out following the protocol based on a previous study (*7*). We prepared double-stranded libraries with the NEBNext Ultra II DNA library preparation kit (New England Biolabs: NEB) following the manufacturer’s instructions, but with minor corrections. Specifically, we diluted the NEBNext Adaptor for Illumina (15 mM) to 1.5 mM with a 10-fold dilution and applied double-sided size selection with 90 µL of Agencourt AMPure XP solution (Beckman Coulter) at both removing long fragments and recovering short fragments. A new blank was processed in parallel with each sample throughout library preparation. To check the percentage of human endogenous DNA and deamination levels, the initial sequencing using MiSeq (Illumina) was conducted at Kitasato University or the University of Tokyo.

For Ikawazu and Kosaku Jomon individuals, Uracil–DNA–glycosylase (UDG) treated DNA libraries were prepared (*83*). Approximately 5 ng of DNA extract was added to a reaction with a total volume of 50 µL, consisting of 5 µl 10X buffer Tango (Thermo), 0.2 µl 2.5 mM dNTP (Thermo), 0.5 µl 100 mM ATP (Thermo), 2.5 µl 10 U/ul T4 polynucleotide kinase (Thermo), 3 µl 1,000 U/µl USER Enzyme (NEB), Ultrapure water (Thermo) up to 50 µL. The mixture was then incubated at 37°C for 3 hours. Subsequently, a blunt-ending step was performed by adding the reaction mix 1 µL of T4 Polymerase (Thermo) to the reaction mix, followed by incubation at 25°C for 15 min and then at 12°C for 5 min. The resulting product was purified using the MinElute kit (QIAGEN) with double elution in 30 µL and 23 µl of EBT buffer. The libraries were prepared using the NEBNext Ultra II DNA library preparation kit.

For some Ikawazu Jomon individuals (IK2010-2, IK2010-3, and IK2013-2), libraries were enriched using an in-solution hybridization protocol targeting approximately 1.2 million nuclear SNPs (1240k SNP set) (*84*). UDG-treated DNA libraries were re-amplified with NEBNext Q5 Master Mix (NEB) and P5 and P7 primers to a concentration of approximately 30 ng/µL. In-solution hybridization was performed using myBaits Human Affinities Complete kit (DAICEL arbor biosciences) with a hybridization temperature set to 60°C and incubated for 20 hours per round.

Libraries without enrichment, referred to as “shotgun,” were sequenced on a NovaSeq6000 (Illumina) at the National Institute of Genetics in Japan, employing 2×100bp paired-end, a NovaSeq6000 at Macrogen in Japan with 2×150bp paired-end, or a HiSeq2500 (Illumina) at Genewiz in Japan with 2×100bp paired-end.

Enriched libraries, designated as “1240K capture,” were sequenced on a MiSeq at the University of Tokyo, employing 2×75bp paired-end.

### Genetic Sex determination

The genetic sex of our samples was determined following the methodology outlined by Skoglund et al., (2013) (*85*). In brief, we computed the ratio of Y chromosome reads to the total number of sex chromosomes (Ry) and classified individuals as females if Ry < 0.016 or males if Ry > 0.075 (refer to table S1 for details).

### Ancient DNA authentication

The authenticity of the samples was assessed through several measures. Firstly, we examined typical patterns of deamination towards read ends and estimated contamination rates on mitochondrial DNA (mtDNA) and Y chromosome in males. Deamination patterns were assessed with mapDamage 2 version 2.2.0 (fig. S1) (*86*).

The NEBNext Ultra DNA Library Prep Kit employs adaptors containing an uracil base, which is cleaved by the USER enzyme before library amplification. However, it’s worth noting that ancient DNA also harbors uracil bases at the 5’ end after the end repair. Consequently, the USER enzyme remove any uracil residuals at the 5’ end, resulting in the absence of deamination damage in that region (*87*).

To evaluate mtDNA contamination, sequence reads were aligned to the revised Cambridge Reference Sequence (rCRS) mitochondrial genome (*88*), and reprocessed the mapped data were processed using the same pipeline as described in the following section. The mtDNA contamination rate was assessed by calculating heterozygosity on the mtDNA using MitoSuite v1.1.0 (*89*). For determining nuclear genome contamination rates in males, heterozygosity was calculated on the X chromosome utilizing ANGSD v0.939 (*90*).

### Sequence data processing and genotype imputation of ancient individuals

In addition to the 25 newly sequenced Jomon individuals, the FASTQ files of previously published Jomon individuals (n = 17) (*7*, *14–16*, *91*) were downloaded from the DNA DataBank of Japan (DDBJ) and the European Nucleotide Archive (ENA). Prior to mapping, AdaptorRemoval v2.3.1 (*92*) was used to trim ambiguous bases at the termini (--trimns), low-quality bases at the termini (--trimqualities), and short reads (--minlength 35), as well as to combine paired-end reads into a consensus sequence (--collapse). The processed reads were then mapped to the human reference genome (hg38) using the BWA MEM algorithm (https://github.com/lh3/bwa). Subsequently, the BAM files generated by BWA MEM were refined with Picard (http://picard.sourceforge.net) CleanSam, and duplicate reads were removed using Picard MarkDuplicates. The ends of the mapped reads were trimmed by TrimBam v1.0.15 in BamUtil (*93*): for non-UDG treated samples, two base pairs (bp) from the 3’ ends and ten bp from the 5’ ends were hard-clipped; for UDG-treated samples, two bp from both the 5’ and 3’ ends of mapped reads were trimmed. After merging BAM files from the same individual into one BAM file, mapped reads with a mapping quality below a Phred score of 30 were filtered out using SAMtools v1.6 (*94*).

We then conducted genotype imputation and haplotype phasing for the Jomon genomes including low-coverage individuals using GLIMPSE2 (*17*). We used phased VCF files from 9,290 modern Japanese individuals, reported by Kawai et al., (2023) (*18*), as the reference panel, along with 2,482 individuals from the 1000 Genomes Project including 104 Japanese individuals (*19*). We followed the tutorial on the GLIMPSE2 website. First, the reference panel was divided into chunks using the GLIMPSE2_chunk command and converted into GLIMPSE2’s binary file format with GLIMPSE2_split_reference. We then ran the GLIMPSE2_phase command for each chunk to perform genotype imputation and phasing, and the imputed chunks of Jomon individuals were ligated using the GLIMPSE2_ligate command. To check the accuracy of the genotype imputation in low-coverage Jomon individual genomes, we downsampled the high coverage data of Jomon individuals (IK2010-1 and FUN23) to low coverage (1x) using samtools and compared the original and imputed genotypes with GIMPSE2_concordance. The imputed genomes of 42 Jomon individuals were combined with the VCF files of 2,482 individuals from the 1000 Genomes Project using the bcftools merge command for subsequent analyses. The 42 Jomon individuals were categorized by their regional origins within the Japanese archipelago, based on their sampling sites, as follows: 2 from Hokkaido, 27 from Kanto, 6 from Tokai, 4 from Hokuriku, and 3 from Chugoku-Shikoku (fig. S2, table S1 and S3).

### Principal component analysis, ADMITURE and Treemix analysis

We carried out principal component analysis (PCA) and ADMIXTURE analysis on the 42 imputed Jomon genomes and 503 East Asian (EAS) genomes from the 1000 Genomes Project, which includes 104 Japanese (JPT), 103 Han Chinese (CHB), 104 Southern Han Chinese (CHS), 93 Dai Chinese (CDX), and 99 Kinh Vietnamese (KHV) genomes. We selected 1,310,568 bi-allelic single nucleotide polymorphisms (SNPs) with a minor allele frequency (MAF) exceeding 35%. The PCA was executed using PLINK v1.9 (*95*), while ADMIXTURE analysis was performed with ADMIXTURE version 1.3.0 (*20*), exploring different values of K (ranging from K=2 to K=5). Cross-validation analysis was conducted, and the configuration of K=3, which exhibited the lowest cross-validation (CV) error, was chosen.

Phylogenetic analysis was performed using TreeMix (*96*) to elucidate the phylogenetic relationships among regional Jomon populations. We analyzed 41 of the 42 Jomon individuals, excluding JpKa6904 from Chugoku-Shikoku (see fig. S2) (*15*) due to its considerable antiquity compared to the other Jomon specimens, which might distort accurate regional phylogenetic relationships of the Jomon people. For comparative analysis, we incorporated 503 genomes from EAS, along with 107 Yoruba (YRI) and residents of Utah (CEPH) of Northern and Western European ancestry (CEU) from the 1000 Genomes Project. From this dataset, we extracted 1,976,831 bi-allelic SNPs with a MAF exceeding 30%. In the analysis, considering the formation of modern Japanese through the admixture of the Jomon people and continental East Asians, we assumed a single admixture pulse from the Jomon lineage to the continental East Asian lineage.

### Estimation of genome-wide genealogy and population size change

To ascertain the population history of the Jomon people (i.e., the effective population size change and the divergence time from other Asians), we constructed genome-wide genealogies of the Jomon people and contemporary East Asians by Relate software (*22*). We used 40 Jomon individuals with available radiocarbon dates and 503 continental East Asian individuals from the 1000 Genomes Project for the analysis.

Input files for Relate were prepared using RelateFileFormats and PrepareInputFiles.sh. Ancestral sequences of *Homo sapiens* (GRCh38) from Ensembl release 103 were utilized for the ancestral sequence analysis, and the StrictMask from the 1000 Genomes Project was used for the genomic masking.

We estimated the genome-wide genealogy with the “Relate” command, setting the mutation rate at 1.25 × 10^-8^ per base per generation (-m 1.25e-8) and an effective population size to 30000 (-N 30000). Sample ages (generations before present) of each Jomon individual were set by --sample_ages option, assuming generation time in humans for 28 years. We then estimated the population size change and divergence times between the Jomon people and the other continental East Asians with the EstimatePopulationSize.sh script, using the estimated genome-wide genealogies of the Jomon people and modern East Asians (.anc, .mut) as input file.

### Archaic introgression

We conducted SPrime analysis (*34*) to identify archaic segments introgressed from Neanderthal or Denisovan into the genomes of Jomon people. SPrime is a reference-free tool for designed to detect such segments by leveraging the strong LD present among archaic variants within the genomes of modern populations. We used SPrime v07Dec18.5e2, with YRI from the 1000 Genomes Project serving as a non-admixed modern population. In addition to the Jomon people, we also identified archaic segments in East Asians from 1000 Genomes Project JPT, CHB, CHS, CDX, and KHV. We adopted the default settings for all SPrime optional parameters, thereby classifying genetic segments with an SPrime score exceeding 100,000 as “archaic segments.” To calculate the match rate to Denisovan (*97*) and Altai Neanderthal (*98*), we followed the same method described in the original paper of SPrime, i.e., the number of matches divided by the number of compared sites for each genetic segments identified as archaic by SPrime. We considered archaic segments with a Match rate to Altai Neanderthal < 0.2 and a Match rate to Altai Denisovan > 0.2 to be introgressed segments from Denisovan lineages. We downloaded the whole genome VCFs of Altai Neanderthal and Denisovan that were mapped to the hg19 reference sequence from http://cdna.eva.mpg.de/neandertal/altai/. To calculate the match rate, the chromosomal positions of the detected Jomon and East Asian archaic segments were converted from hg38 to hg19 using liftOver (*99*).

### Genome-wide selection scan

To investigate the genetic adaptations of the Jomon lineage after their early divergence from the continental East Asians, we detected positive natural selection signals. Given our interest in discerning differences in genetic adaptation between the Jomon people and continental East Asians, we employed the cross population nSL (XP-nSL) method to detect regions with particularly long-range haplotypes (i.e., selective sweep signals) in the Jomon people.

XP-nSLs were calculated for 24,704,448 SNPs by Selscan software (*45*). A significantly positive XP-nSL value suggests signals of positive natural selection in the Jomon lineage, whereas a significantly negative value indicates positive natural selection in the Han Chinese.

To identify phenotypes that spread among the Jomon people through positive natural selection, we devised the regional XP-nSL (rXP-nSL) index by combining XP-nSL with summary statistics (β and *P-*values) obtained from previous genome-wide association studies (GWAS) on 62 traits in modern Japanese (*46–49*). Given that the GWAS data is based on the hg19 sequence, we converted chromosomal positions from hg19 to hg38 using liftOver.

The rXP-nSL index is designed to detect trait-associated genetic regions enriched with SNPs exhibiting high XP-nSL values. In this study, we utilized a technique known as linkage disequilibrium clumping (LD-clumping) to precisely define “trait-associated genetic regions,” ensuring that they are unequivocally independent of each other. Although the goal of GWAS is to identify specific SNPs that affect diseases or traits, SNPs that appear to be associated due to LD can also be detected around SNPs with true effects. LD-clumping integrates results from GWAS with LD information to select a representative SNP from each cluster of trait-associated SNPs with LD. In this study, ’trait-associated genetic regions’ were identified as clusters of trait-associated SNPs that are in linkage disequilibrium (LD) within 1MB window, using LD-clumping.

We combined GWAS data of modern Japanese individuals (effect allele, effect size β, and P-value for each SNP) from previous studies with VCF files of the modern Japanese (JPT) from the 1000 Genomes Project. Using PLINK, we executed LD-clumping with the options --clump-p1 5e-8, --clump-kb 500 --clump-r2 0 for 1KG JPT modern Japanese genomes. We adopted --clump-kb 500 to set the window length at 1MB. With --clump-p1 5e-8, we set the GWAS P-value threshold at 5.0 × 10^-8^, and with --clump-r2 0, we set the linkage disequilibrium coefficient r^2^ to be greater than 0, considering trait-associated SNP pairs that are within 1MB window as being in the “same” trait-associated region. Subsequently, we calculated the mean value of XP-nSL at the *n* trait-associated SNPs within the trait-associated region of the Jomon people (fig. S7). The rXP-nSL was then calculated by the following procedure: (1) sample a 1MB region in the whole genome; (2) randomly select the same numbers of SNPs (*n* SNPs) as the target trait-associated region from the sampled 1MB region and obtain their mean XP-nSL; (3) repeat (1) and (2) 100 times to obtain the null distribution of the mean XP-nSL for 1MB trait-associated regions with *n* SNPs; (4) Based on the mean and standard deviation of XP-nSL distribution, obtain rXP-nSL by the following formula;

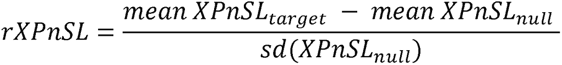

Trait-associated genetic regions with rXP-nSL greater than 6.566567, which is the 95-percentile of 1,481 genetic regions in total among 62 trait GWASs, were interpreted as having undergone positive natural selection in the Jomon lineage (fig. S8). For each trait-associated genetic region, we evaluated the direction of the effect of the haplotype that underwent natural selection in the Jomon lineage by comparing the mean 2βf within each trait-associated genetic region, the mean value of the product of 2 times the effect allele frequency of each SNP and the effect size (β) in the focal trait-associated genetic region, between the Jomon people and the Han Chinese. If the mean of 2βf was greater for the Jomon people than for the Han Chinese, we considered it as having a “relatively positive effect” on the Jomon people. Conversely, if the mean of 2βf was smaller for the Jomon people than for the Han Chinese, we regarded it as having a “relatively negative effect” on the Jomon people.

### Estimation of allele frequency trajectory

To elucidate the timing of positive natural selection within the population history of the Jomon people, we conducted CLUES2 (*50*, *51*) analysis to estimate the allele frequency trajectories and selection coefficients of trait-associated SNPs which exhibited the significant selective sweep signals in the Jomon lineage. CLUES2 can be run on ancient genomes which makes use of ancient genotype probabilities, sample ages, the Relate output genealogies (.anc, .mut) and the population size change (.coal). In this study, we selected the following SNPs as representative within two trait-associated regions where signals of positive natural selection were observed, based on their having the lowest GWAS P-values: rs1162015 C/T, a derived allele on the *FTO* gene on chromosome 15 associated with increased BMI (*P-*value = 4.9 × 10^-72^), and rs662799 A/G, a derived allele on the *APOA5* gene on chromosome 11 associated with increased triglycerides (*P-*value < 1 × 10^-320^) (*48*). We adhered to the procedure outlined in the CLUES2 tutorial as follows: 1. sample the gene trees for the focal SNPs by the sampleBranchLengths.sh script of Relate package; 2. convert output trees (.newick) to the CLUES2 input file (_times.txt) by the RelatetoCLUES.py command; 3. estimate the allele frequency trajectory by CLUES2’s inference.py command, using coalescent times (_time.txt), genotype probabilities and sample ages of the Jomon individuals (AncientSamples file) and population size change through time (.coal). However, when applying these steps to our sample, we encountered a unique challenge not directly addressed by the standard CLUES2 procedures. Our analysis, which is based solely on ancient genomes, revealed a limitation related to the coalescence rate estimated for periods newer than 96 generations ago (with IK002 (*7*, *91*) being the most recent of our Jomon samples) being zero, which led to errors in running the python inference.py command with the *.coal files for population size change in the Jomon people. Given the distinct evolutionary history of the Jomon lineage, incorporating modern samples into the analysis to generate the .coal file seems inappropriate for our study. To address the zero coalescence rate problem, we contemplated adjusting the timeframes of all our ancient samples and the .coal file to make the most recent sample’s generation (96 generations ago) corresponds to time 0 (the most recent epoch). Specifically, we manually edited the AncientSamples file by subtracting 96 from the generation count for each sample, thereby setting 96 generations ago as the new time 0. For the .coal file, we removed the coalescence rate data for generations 0-68.9535 and sliding the remaining coalescence rate and generation times towards the newer side, as illustrated below:

Original:

0

0 35.7143 49.6248 68.9535 95.8106 133.128 184.981 257.031 357.143 496.248 …

#generation times

0 0 0 0 0 0 9.72E-05 8.27E-05 0.000127535 0.00016142 … #coalescence rates

Adjusted:

0

0 37.3174 51.853 72.05 100.112 139.105 193.287 268.571 373.174 518.53 …

#modified generation times

0 0 9.72E-05 8.27E-05 0.000127535 0.00016142 0.00019344 0.000198509

0.000179675 0.000146651 … #modified coalescence rates

Therefore, we need to emphasize that the allele frequency trajectories for the Jomon people estimated in our study started from 96 generations ago, which is the newest sample age of our Jomon individuals. We estimated the selection coefficient, s, by dividing ages into periods during and after the Last Glacial period, using the --timeBins option with a cutoff at 10,000 years ago.

### Comparison between the Jomon people and Greenlandic Inuit for frequency of alleles associated with diet and cold adaptation

Previous studies on the population genomics of Greenland Inuit (GI) have detected signals of positive natural selection at various loci, such as *FASD1, FADS2*, and *FADS3* on chromosome 11, *TBX15* and *WARS2* on chromosome 1, and *FN3KRP* on chromosome 17 (*65*). These loci are thought to be associated with adaptations to the cold climates of their settlement areas and their diets rich in omega-3 polyunsaturated fatty acids (PUFAs), which are prevalent in seafood (*65*).

Considering the potential similarity in cold environment adaptations between the Jomon and Greenlandic Inuit, it is plausible that the allele frequencies in the Jomon people exhibiting positive natural selection signals observed in Greenlandic Inuit would more closely resemble those of Greenlandic Inuit than populations sharing a more recent common ancestor with the Jomon people, such as Han Chinese. To explore this hypothesis, we calculated the *f4(CEU, GI; Jomon, CHB)* (*100*) statistic for genome-wide SNPs.

Negative *f4* values indicate that allele frequencies in the Jomon people are closer to those in Greenlandic Inuit than to those in Han Chinese. We obtained the allele frequencies in Greenlandic Inuit for 94,577 genome-wide SNPs from the previous study (*65*) and performed the *f4(CEU, GI; Jomon, CHB)* calculation using a custom script in R.

## Acknowledgments

We thank Dr. Kazuki Morisaki at the University of Tokyo for his advice on the Upper Paleolithic of Paleo-Honshu Island. We also thank Dr Yosuke Kaifu in University of Tokyo, and Dr Catherine Walker in University College London, for helpful discussion. Computations were partially performed on the NIG supercomputer at ROIS National Institute of Genetics.

## Funding

JSPS KAKENHI Grant Number 23H04836 (YY, MI, HO)

JSPS KAKENHI Grant Number 23H04838 (HO, YUW)

JSPS KAKENHI Grant Number 23H04840 (MI, JO)

JSPS KAKENHI Grant Number 23KJ0654 (YOW)

JSPS KAKENHI Grant Number 22H00020 (RT, MY, HO)

JSPS KAKENHI Grant Number 22H00718 (YY, MY, SM, HO)

JSPS KAKENHI Grant Number 22H00421 (JO)

JSPS KAKENHI Grant Number 21H04779 (HO, JO, DW, TG, YUW)

JSPS KAKENHI Grant Number 21H00337 (HO)

JSPS KAKENHI Grant Number 22KF0092 (HO)

JSPS KAKENHI Grant Number 21H05362 (HO)

JSPS KAKENHI Grant Number 21K15175 (YUW)

JSPS KAKENHI Grant Number 21K19289 (HO)

JSPS KAKENHI Grant Number 21K19289 (HO)

JSPS KAKENHI Grant Number 18H03590 (RT, MY, GV, HO)

JSPS KAKENHI Grant Number 18H03593 (YY, MY, TG, SM, HO)

JSPS KAKENHI Grant Number 17H01453 (TN, TK, HO)

JSPS KAKENHI Grant Number 17H03738 (HO, TK)

JSPS KAKENHI Grant Number 26251050 (HO)

JSPS KAKENHI Grant Number 25284157 (YY, MY, SM, HO)

JSPS KAKENHI Grant Number 16H06279 (PAGS) (AT, HO)

JSPS KAKENHI Grant Number 22H04925 (PAGS) (AT, HO)

THE MITSUBISHI FOUNDATION 202110009 (HO)

## Author contributions

Conceptualization: YUW, HO

Methodology: YUW, JO

Investigation: YUW, YOW, DW, GV, AKT, YN, TS, ATT, SM, TON, MY, TM, NCBN

Visualization: YUW, YOW, TS

Funding acquisition: HO, MY, YY, RT, YUW, YOW

Project administration: HO

Supervision: KK, TG, TK, MO, KH, MY, YY, RT, JO, HO

Writing – original draft: YUW, YOW, TAN, MI, HO

## Data and materials availability

All raw genomic data (fastq files) are available for download in the DNA DataBank of Japan (DDBJ) Sequence Read Archive (DRA. https://www.ddbj.nig.ac.jp/index-e.html) under the accession numbers PRJDB14637, PRJDB18003 and PRJDB18005.

