## Supplementary for "Cold adaptation in Upper Paleolithic hunter-gatherers of eastern Eurasia"

Supplementary Text

**NCBN Controls WGS Consortium**

Hatsue Ishibashi-Ueda,^1^ Tsutomu Tomita,^1^ Michio Noguchi,^1^ Ayako Takahashi,^1^

Yu-ichi Goto,^2^ Sumiko Yoshida,^3^ Kotaro Hattori,3 Ryo Matsumura,^3^ Aritoshi Iida,^4^

Yutaka Maruoka,^5^ Hiroyuki Gatanaga,^6^ Akihiko Shimomura,5 Masaya Sugiyama,^7^ Satoshi Suzuki,^5^, Kengo Miyo,^8^

Yoichi Matsubara,^9^ Akihiro Umezawa,^10^ Kenichiro Hata,^11^ Tadashi Kaname,^12^

Kouichi Ozaki,^13^ Haruhiko Tokuda,^13^ Hiroshi Watanabe,^13^ Shumpei Niida,^13^

Eisei Noiri,^14^ Koji Kitajima,14 Yosuke Omae,^14,15^ Reiko Miyahara,^14^ Hideyuki Shimanuki,^14^ Yosuke Kawai,15 and Katsushi Tokunaga^14,15^

1 NCVC Biobank, National Cerebral and Cardiovascular Center, Suita, Osaka 564-8565, Japan

2 Medical Genome Center, National Center of Neurology and Psychiatry, Kodaira, Tokyo 187-8551, Japan

3 Department of Bioresources, Medical Genome Center, National Center of Neurology and Psychiatry, Kodaira, Tokyo 187-8551, Japan

4 Department of Clinical Genome Analysis, Medical Genome Center, National Center of Neurology and Psychiatry, Kodaira, Tokyo 187-8551, Japan

5 NCGM Biobank, National Center for Global Health and Medicine, Shinjuku-ku, Tokyo 162-8655, Japan

6 AIDS Clinical Center, National Center for Global Health and Medicine, Shinjuku-ku, Tokyo 162-8655, Japan

7 Department of Viral Pathogenesis and Controls, Research Institute, National Center for Global Health and Medicine, Ichikawa, Chiba 272-8516, Japan

8 Center for Medical Informatics and Intelligence, National Center for Global Health and Medicine, Shinjuku-ku, Tokyo 162-8655, Japan

9 National Center for Child Health and Development, Setagaya-ku, Tokyo 157-8535, Japan

10 Center for Regenerative Medicine, National Center for Child Health and Development, Setagaya-ku, Tokyo 157-8535, Japan

11 Department of Maternal-Fetal Biology, National Center for Child Health and Development, Setagaya-ku, Tokyo 157-8535, Japan

12 Department of Genome Medicine, National Center for Child Health and Development, Setagaya-ku, Tokyo 157-8535, Japan

13 Research Institute, National Center for Geriatrics and Gerontology, Obu, Aichi 474-8511, Japan

14 Central Biobank, National Center Biobank Network, Shinjuku-ku, Tokyo 162-8655, Japan

15 Genome Medical Science Project (Toyama), Research Institute, National Center for Global Health and Medicine, Shinjuku-ku, Tokyo 162-8655, Japan

**Treemix analysis**

The 42 Jomon genomes, collected from various regions of the Japanese archipelago as depicted in fig. S2, facilitated our phylogenetic analysis aimed at uncovering the regional genetic diversity of the Jomon people. According to Jeong et al., (2023), JpKa6904, the earliest Jomon specimens sequenced to date (*25*), was reported to be an outgroup to the more recent Jomon individuals by f-statistic based approaches (*49*). Our phylogenic analysis, excluding JpKa6904 individual, showed that the northwestern Jomon lineage diverged from the southwestern Jomon lineage, suggesting a gradient of genetic diversity that extends from the southwest to the northeast across the Jomon population. The 41 Jomon individuals examined in this study span a broad temporal range, from approximately 6,000 to 2,700 years ago in each geographic region in the Japanese archipelago. Consequently, our results likely reflect geographical diversity within the Jomon population rather than temporal variations.

**The correlation between our findings and the archaeological evidence from the Upper Paleolithic (UP) in the Japanese archipelago**

The UP occupation began around 38,000 yr cal BP (*50*) and continued until about 15,000 yr cal BP on P-Honshu, extending from 15,000 to 12,000 yr cal BP on Hokkaido, south of PSHK (*51*, *52*). The UP period is divided into three sub-stages – early, middle, and late – characterized by lithic reduction patterns, toolkit compositions, stone-tool types, and date of the sites. Notably, the middle UP stage, spanning approximately 30,000-20,000 yr cal BP, shows heightened mobility with a blade-based tool-kit assemblage (*51*, *53*–*55*).

During the middle UP, the blade-point industry predominantly flourished particularly for the northern P-Honshu, marked by prismatic and conical blade cores, and by relatively large blade-based tools including backed points, basally retouched points, burins, endscrapers and sidescrapers, exhibiting a composition akin to UP assemblages found elsewhere in northern latitude of Asia. These tools were highly standardized with intensive retouch, and toolkit diversity became clearly higher than the small flake-based trapezoid industry during the early UP (*53*, *56*). This particular blade-based reduction strategy was facilitated by the availability of high-quality raw materials such as obsidian and siliceous shale. Tools and blades were mostly manufactured at a limited number of hub sites nearby high-quality raw material sources, and transported to distant logistic satellite sites, obviously suggested that more scheduled foraging activities accomplished than the previous early UP assemblages. This period witnessed increased repeated occupation at certain locations and reduced residential movements, augmented by short-term, task-specific extraction locations (*53*, *57*).

Following the initial appearance of modern human at 38,000 yr cal BP, drastic change in the chronological distribution of radiocarbon dates were observed by the Kernel Density Estimate model (*8*, *58*, *59*) around 30,000 yr cal BP, both on P-Honshu and PSHK during Middle UP(*60*). These technological changes and the fluctuation of cultural occupation patterns align with the hypothesis of the population replacement(s) occurred between the early and middle UP periods on P-Honshu and PSHK. Moreover, they suggest that the middle UP foragers adopted to the colder landscape of the northern Asia before colonized into P-Honshu.

Given the changes observed in cultural evidence during the middle UP, coupled with the estimated divergence time between the Jomon and continental East Asians (Fig. 3 and fig. S2), it is possible that the isolation of the Japanese archipelago from the East Eurasian continent just before or during the LGM resulted in a divergence between the UP populations on the Eurasian continent and those on the Japanese archipelago, leading to cultural transitions. The ecological background underwent drastically alteration from the temperate Pan-mixed forest dominated landscape, characterized by *Palaeoloxodon-Shinomegaceroides* complex by 30,000 yr cal BP, to a cool-temperate coniferous forest with patchy grassland accompanied by Mammoth fauna in the PSHK during the LGM, showing different pattern of smaller regional biomes (Fig. 1) (*53*, *61*, *62*). The colder environment of both P-Honshu and PSHK during LGM shared several common faunal and floral species with those of continental northeastern Asia, possibly supporting migration from the northern landscape.

The subsequent decline in the frequency of cultural occupation preceding to the Jomon period (*60*) closely aligns with our finding (Fig. 3), indicating a rapid decline in population size until the beginning of the Jomon period. The period around 20,000 years ago, encompassing the LGM, likely presented significant challenges for the UP population in the Japanese archipelago, potentially leading to a substantial decline in population size. Following the massive increase in effective population size after the LGM (Fig. 3), it is likely that the UP population, survivors of the ice age, formed the Jomon culture in the warming Japanese archipelago, epitomized by its adoption of distinctive pottery (*63*). The divergence between the Hokkaido Jomon and the Hondo Jomon occurred between 8,800 and 10,000 years ago. Archaeologically, it has been suggested that the human population in P-Honshu migrated to PSHK around 13,000 years ago (*60*), a finding supported by Fig. 3B, figs. S2 and S4 (*52*). Moreover, the absence of recovery in the effective population size in the Hokkaido Jomon lineage is consistent with the line of cultural evidence (*53*, *56*, *64*, *65*), indicating a decrease in the intensity of cultural occupation in Hokkaido around 15,000 years ago.

**Our findings and their implications for the initial peopling of the Americas**

We have confirmed that the Hokkaido Jomon peoples, represented by Funadomari individuals, were diverged from a single lineage sharing with the Honshu Jomon peoples (Fig 3 and fig. S2). All Jomon individuals examined so far thus has consistently indicated a shared common ancestor. These findings from our analysis may offer new insights into the initial peopling of the Americas.

The discovery of bifacial stemmed points at the Cooper’s Ferry site (Idaho, USA), dating back to ~15,780 yr cal BP, indicates a closest similarity to Late UP stemmed point technology in PSHK. This suggests the existence of cultural connections with UP peoples at the Pacific Rim, potentially tracing back to some of the earliest peoples in the Americas between ~22,000 and 16,000 years ago (*66*, *67*). Other archaeological evidence also supports this hypothesis; for instance, bifacial stemmed point technology of this old did not emerge in any region of northeast Asia, including Beringia until around 14,200 yr cal BP (*68*). Despite the cultural evidence supporting this scenario, further discussion must be postponed due to the absence of human skeletal remains and subsequent lack of ancient genomic data from PSHK.

Furthermore, the “Out-of-Japan” hypothesis is challenged by various lines of evidence from bioanthropology. Morphological analysis of skeletal remains, along with data on haploid markers (both on mtDNA and Y-chromosomes), and whole genomic analyses of Holocene Jomon, all suggested that the Jomon peoples are unlikely to be direct ancestors of the ancestral and modern Native Americans (*69*).

However, our analysis indicates that Hokkaido Jomon peoples diverged from those of Honshu Jomon around 10,000-8,000 years ago (Fig. 3 and fig. S4). The UP peoples who inhabited PSHK before the migration of Jomon peoples to the isolated island of Hokkaido in Holocene likely belong to a different lineage, diverging from ancestral East Eurasians. As previously suggested, UP in PSHK, dated between ~30,000 and 12,000 yr cal BP, exhibited a higher diversity of lithic assemblage, including various stone tool technologies and lithic reduction sequences such as small flake-based, blade-based, flake-based, bifacial, and microblade (*51*, *56*, *70*). This higher diversity in lithic assemblage in PSHK may suggest a more complex prehistory of human migrations in and out of the region, situated in the ecotone between the cool-temperate coniferous forest to the south and patches of sub-arctic open forest and steppe to the north (*60*, *65*). These pre-Jomon peoples in PSHK still stand as candidates for the first Americans. Further testing of the “Out-of-Japan” hypothesis or coastal migration theory is challenging based solely on the Jomon genomes presented in this study. Detailed analysis of the UP peoples in PSHK is imperative.

**Archaic segments**

We initiated a quest to identify archaic segments using the imputed genomes of 40 Jomon individuals employing SPrime (*71*), juxtaposed with data from continental East Asians sourced from the 1KG (see fig. S5). Previous studies have discussed multiple admixture events between Denisovans and the East Eurasians, based on the distribution of match rate to Altai Denisovan genome (*71*, *72*). Our analysis of Jomon individuals revealed 950 archaic-like segments scattered across the genome, whereas a mean of 1930 such segments were detected in five East Asian populations. Detecting low-frequency archaic segments in imputation data is fraught with challenges due to low imputation quality (Fig. 2A). Consequently, whole genome sequencing data is expected to yield a larger number of detectable archaic segments than imputed genomes. Therefore, rather than a quantitative comparison for Denisovan segments, our focus shifted to examining differences in the distribution of Match to Altai Denisovan among populations, aiming to determine whether the distribution of archaic segments differs between the Jomon people and other continental East Asians. Our analysis revealed no significant differences in the distribution of archaic segments between them (fig. S5A and S5B). The absence of difference in archaic segments between the Jomon and other East Eurasians suggests that admixture events with Denisovans occurred prior to the divergence of the Jomon people from continental East Asian groups, namely, before 27,000 and 19,000 years ago (Fig. 3 and fig. S3). It is probable that there exist Denisovan-derived segments, yet to be discovered, which are scarcely transmitted to modern Japanese individuals, remaining at very low frequencies among contemporary individuals, thereby posing challenges for imputation methodology.

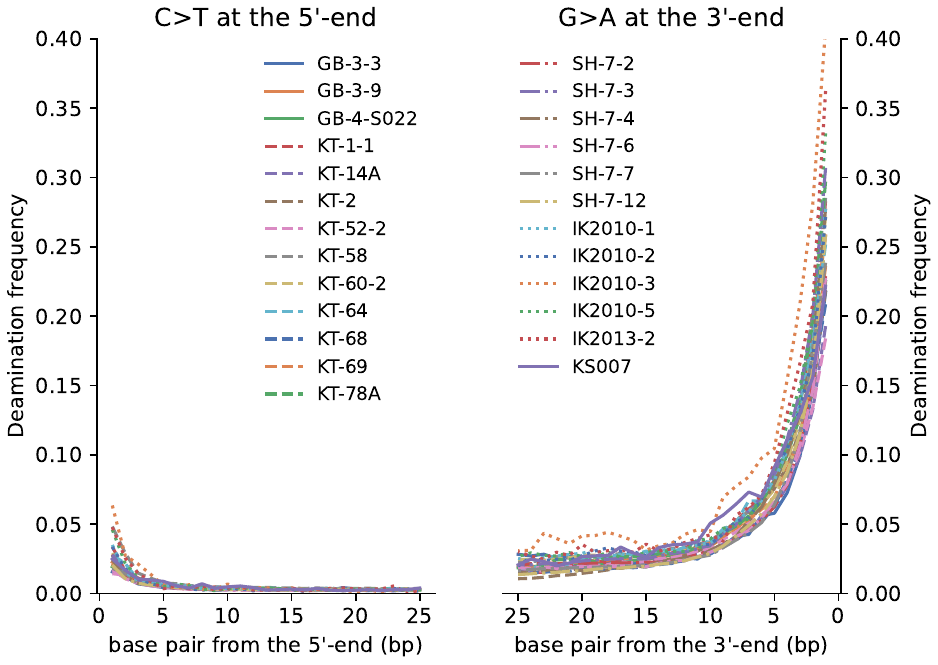

**Fig. S1. DNA damage plots for newly sequenced Jomon individuals.** Damage patterns on the left plot show C>T misincorporations at the 5' end, while those on the right plots show G>A misincorporations at the 3' end.

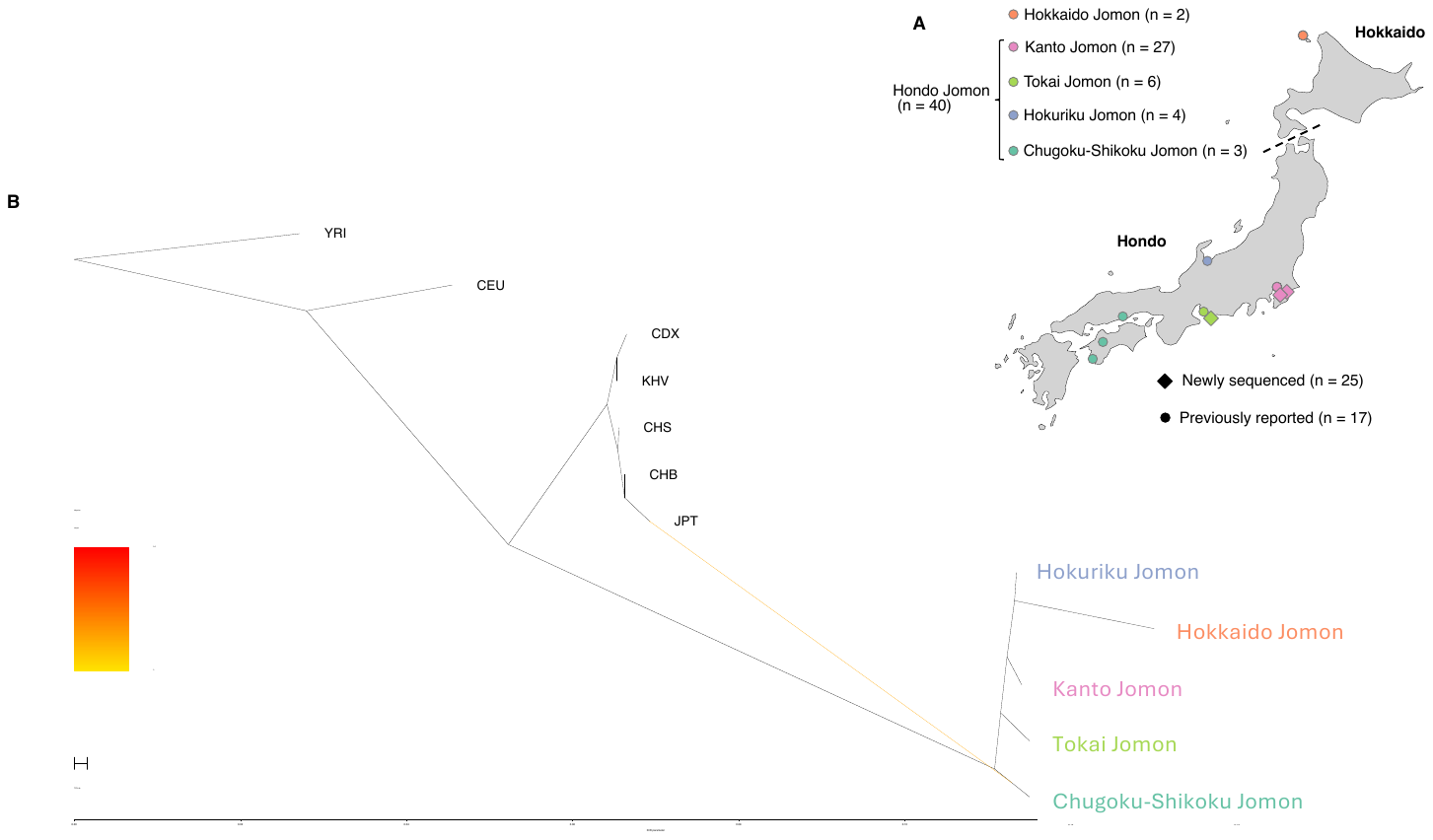

Fig. S2. Sampling locations and phylogenetic analysis of the Jomon people. (A) Archaeological sites are marked with diamonds for newly sequenced samples in this study and circles for previously reported samples. The colors represent regions in the Japanese archipelago. In this study, regions except for Hokkaido are collectively referred to as “Hondo”. The current Hondo region was a single island known as the Paleo-Honshu Island during the Last Glacial Maximum (LGM) period, due to the lowering of sea levels, while Hokkaido was connected to the continental Asia and known as the Paleo-Sakhalin Hokkaido Kurile Peninsula (PSHK) (Fig. 1). (B) Maximum likelihood phylogenetic tree of regional populations of the Jomon people reconstructed by *TreeMix* under a model of one migration.

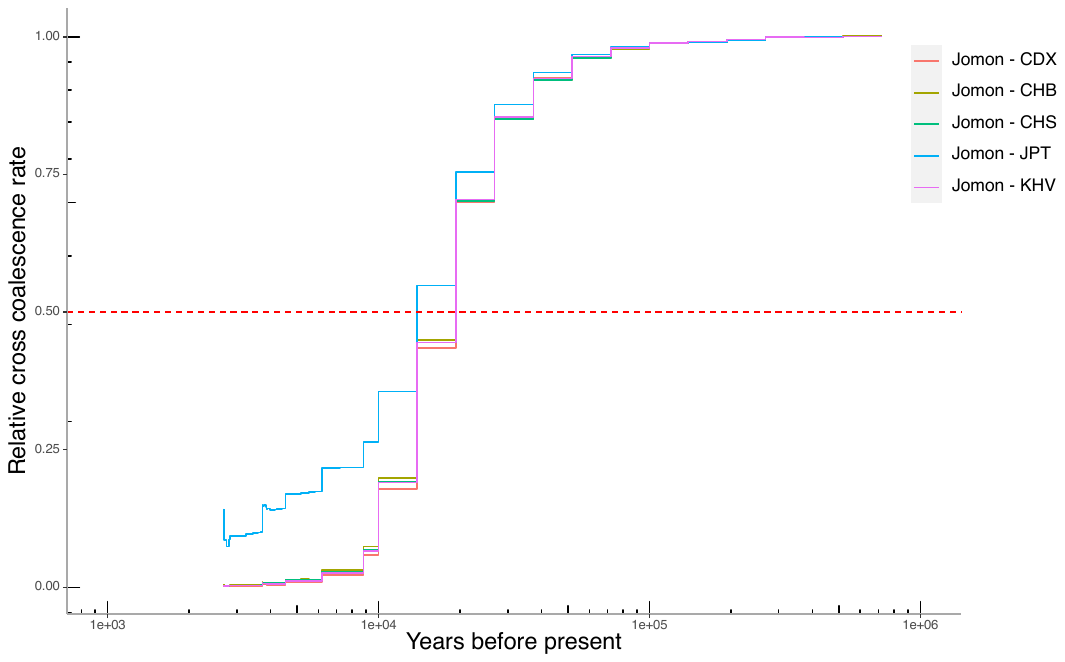

Fig. S3. Relative cross coalescence rate (RCCR) between the Jomon people and modern 1KG East Asians. RCCR is calculated between pairs of populations, where a generation time at an RCCR of 0.5 indicates the divergence time between the two focal populations.

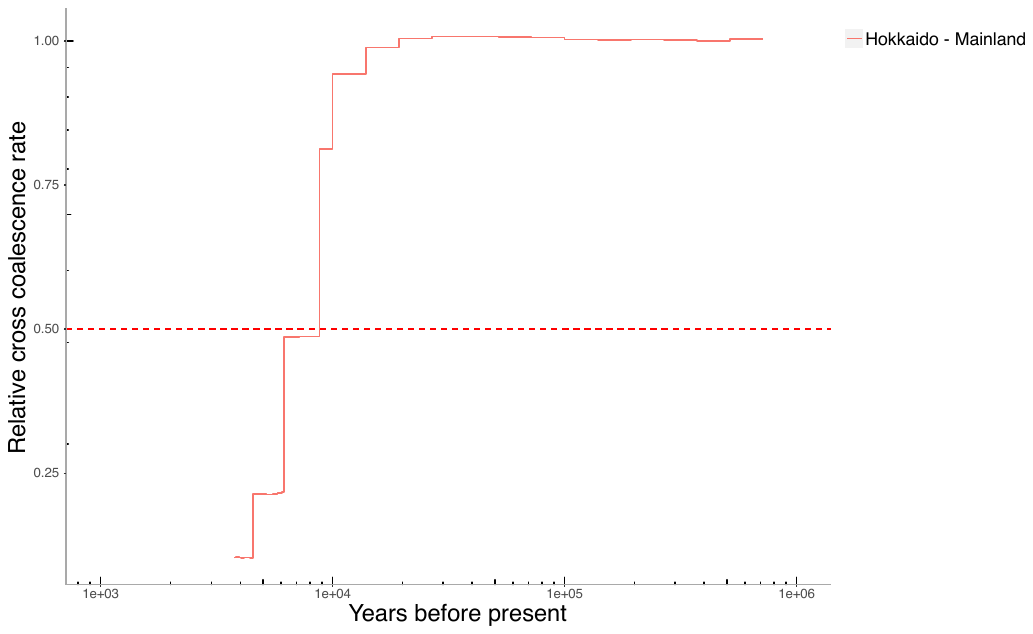

**Fig. S4. Relative cross coalescence rate (RCCR) between the Hondo and Hokkaido Jomon people.**

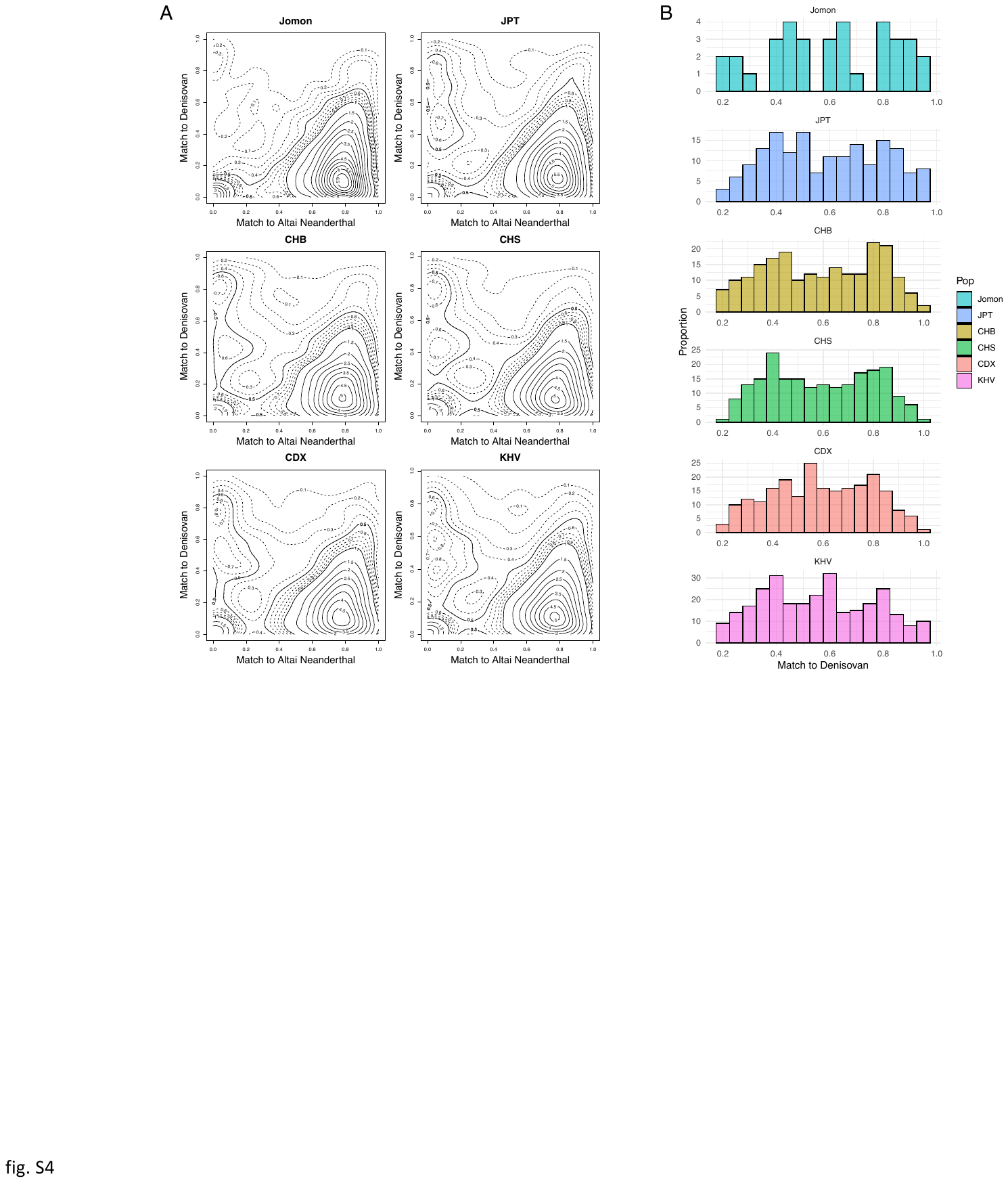

**Fig. S5. Distribution of match to Denisovan/Neanderthal for detected archaic segments in the Jomon individuals and East Eurasians.** We inferred archaic segments of 40 Jomon individuals and 1KG East Eurasians (JPT, CHB, CHS, CDX, KHV) by SPrime software, a reference-free detection method, using 1KG Yoruba (YRI) individuals as a non-admixed population. The match rate is calculated by the number of matches to the given archaic genome divided by the number of putative archaic alleles in the focal segment. (A) Contour density plots of match rate of archaic segments to the Altai Neanderthal/Denisovan. (B) Histograms of Match to Denisovan for the archaic segments with Match to Neanderthal < 0.2 and Match to Denisovan > 0.2, which we deemed as derived from Denisovan lineage. In previous studies, the non-unimodal distributions of Match to Denisovan were considered to indicate admixture events with multiple Denisovan lineages (*36*, *72*).

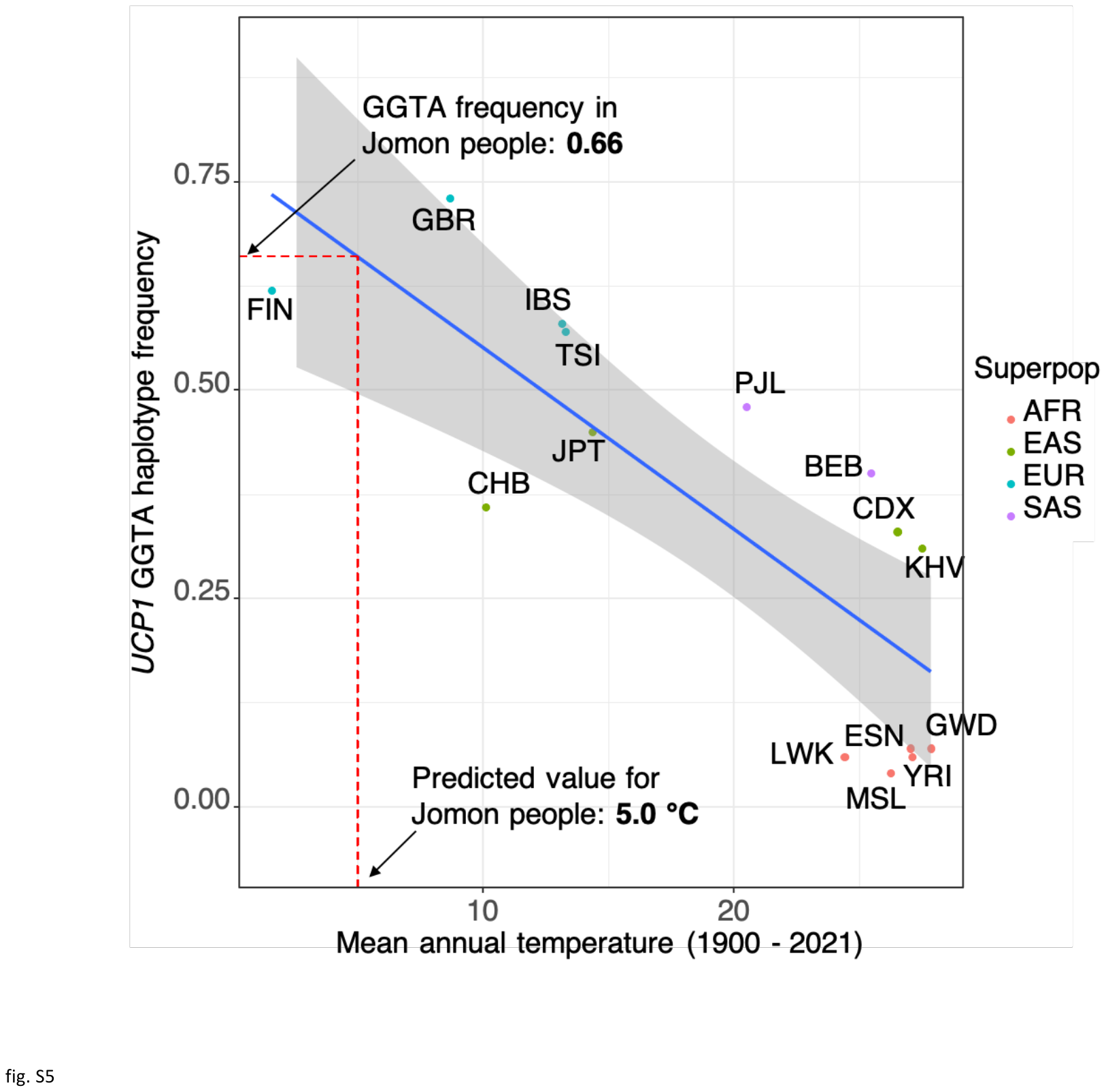

**Fig. S6. Correlation between the *UCP1* GGTA haplotype frequencies and mean annual temperatures of global human populations in the 1KG and the Jomon people**. Blue line and grey band represent a regression line and its 95% confidence interval, which were computed with exception of the Jomon people. The abbreviations used are the following: African (AFR): ESN (Esan in Nigeria), GWD (Gambian in Western Divisions in the Gambia), LWK (Luhya in Webuye, Kenya), MSL (Mende in Sierra Leone), YRI (Yoruba in Ibadan, Nigeria); European (EUR): FIN (Finnish in Finland), GBR (British in England and Scotland), IBS(Iberian Population in Spain), TSI (Toscani in Italia); East Asian (EAS): CDX (Chinese Dai in Xishuangbanna, China), CHB (Han Chinese in Bejing, China), JPT (Japanese in Tokyo, Japan), KHV (Kinh in Ho Chi Minh City, Vietnam); South Asian (SAS): BEB (Bengali from Bangladesh), PJL (Punjabi from Lahore, Pakistan).

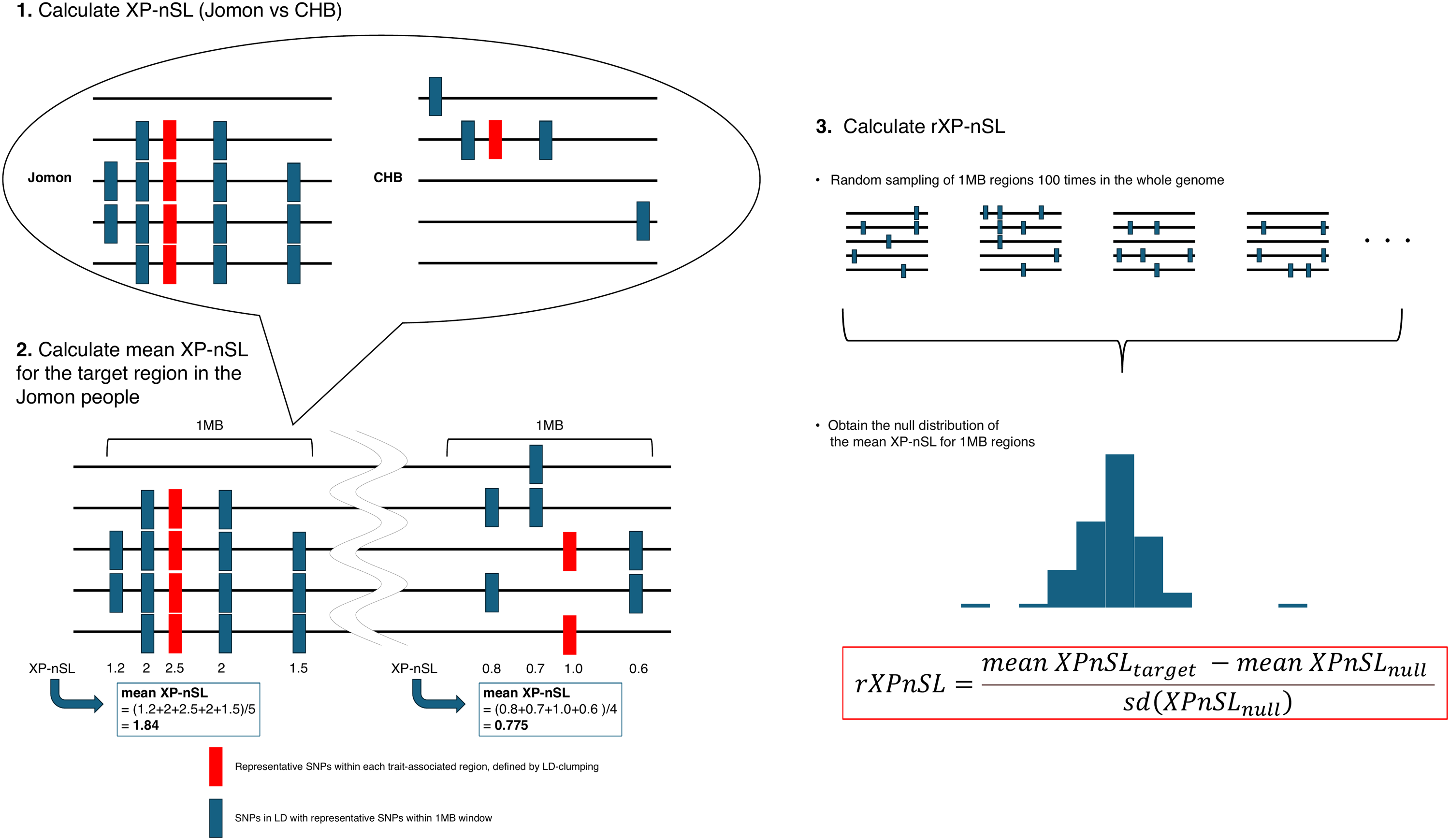

**Fig. S7. Overview of genome-wide selection scan methods of this study.** We executed LD-clumping for 1KG JPT modern Japanese genomes to determine the representative SNP (red) and other trait-associated SNPs (blue) of each trait-associated region.

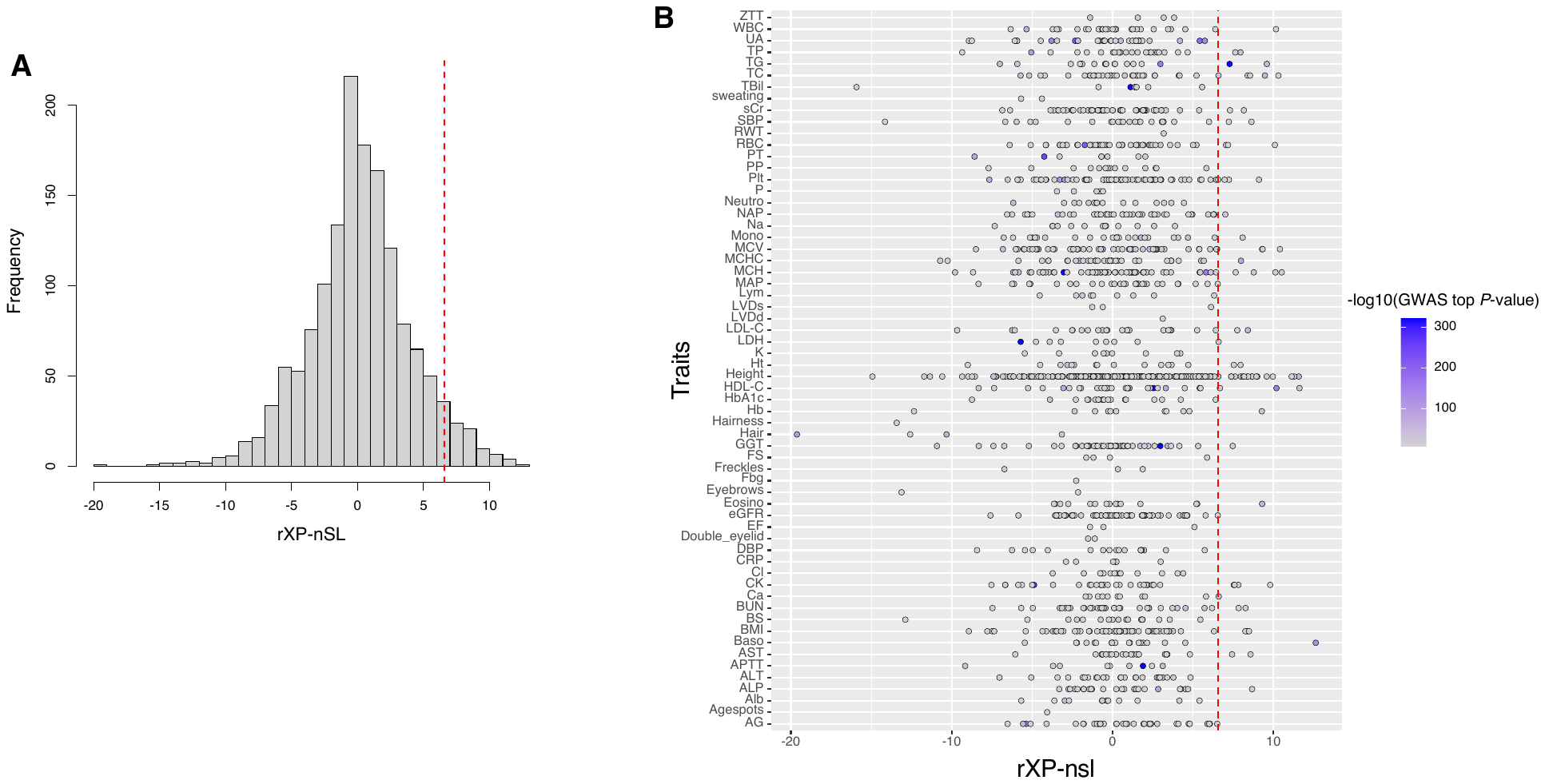

**Fig. S8. rXP-nSL values for trait-associated regions which were detected by previous genome-wide association studies (GWAS).** (A) Histogram of rXP-nSL values of 1481 trait-associated regions. The 95th percentile of the empirical distribution are shown as red dashed vertical line. (B) rXP-nSL values for trait-associated regions of each trait. The vertical axis represents traits cited from GWAS targeting modern Japanese individuals in previous researches (*41*–*44*). Each point represents a trait-associated region. The color of each point is displayed darker for lower P-values in the previous GWAS. The 95th percentile of the empirical distribution are shown as red dashed vertical line.

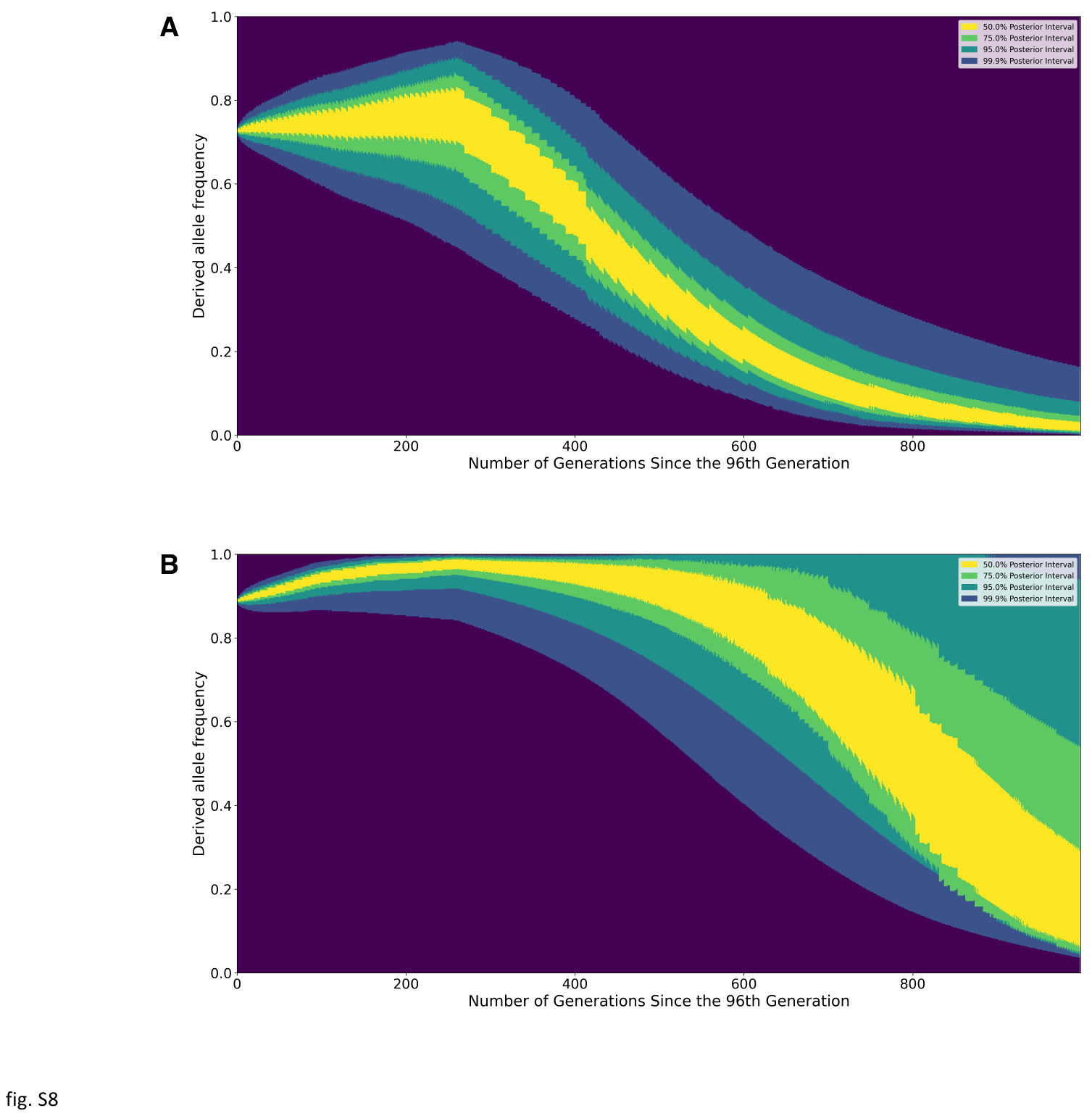

**Fig. S9.** CLUES2 plot for the Jomon people which show the posterior probability of the derived allele frequency trajectory for (A) rs662799 and (B) rs11642015. The derived allele for rs662799 in the *ZPR1-APOA5* region elevates TG, whereas the derived allele for rs11642015 elevates BMI.

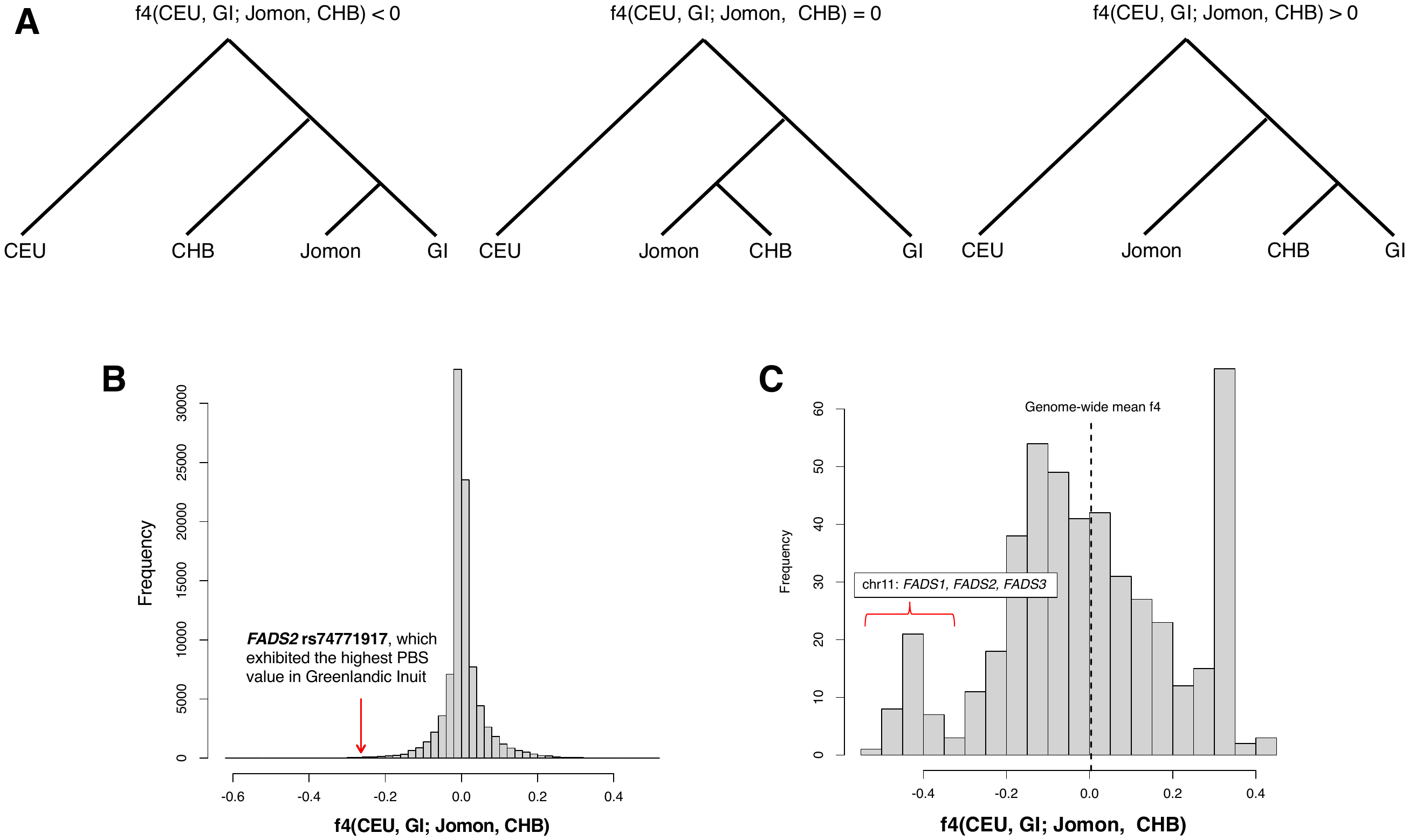

**Fig. S10. Histogram of *f4(CEU, Greenlandic Inuit; Jomon, CHB)* for SNPs with significant PBS values in Greenlandic Inuit (GI) in Fumagalli et al., (2015).** We conducted the *f4* test to clarify whether the allele frequencies in the Jomon people are similar to those in the GI, regarding SNPs with positive natural selection signals in the GI (*47*). (A) Supported topologies of the Jomon people, GI and CHB for various *f4* values. (B) Histogram of *f4(CEU, GI; Jomon, CHB)* for 94,577 whole genome SNPs. The red arrow shows the *f4(CEU, GI; Jomon, CHB)* value of *FADS2* rs74771917, which exhibited the highest PBS value in GI. (C) Histogram of *f4(CEU, GI; Jomon, CHB)* for SNPs with significant PBS values higher than the 99.5th percentile in GI, as reported in the previous study. The mean *f4* value for genome-wide SNPs is 0, indicating that the central topology in (A) was supported across the whole genome. However, for SNPs on the genes *FADS1, FADS2, FADS3* on chromosome 11, which are known to have undergone positive natural selection in the GI (*47*), the *f4* showed significantly negative values, suggesting that the allele frequencies of these SNPs in the Jomon people were found to be significantly closer to those of GI than to Han Chinse, who shares a more recent common ancestor with the Jomon lineage.

Table S1. Newly sequenced sample information.

Table S2. Newly sequenced library information with accession numbers.

**Table S3. List of the Jomon individuals from previous studies used in this study.**

**Table S4. Allele frequencies, PBS in Greenlandic Inuit (GI) and *f4(CEU, GI; Jomon, CHB)* for SNPs with significant PBS values higher than the 99.5th percentile in GI, as reported in Fumagalli et al., (2015).** Allele frequencies in GI and PBS values were cited from the previous study (*47*).

31.	A. Auton, G. R. Abecasis, D. M. Altshuler, R. M. Durbin, G. R. Abecasis, D. R. Bentley, A. Chakravarti, A. G. Clark, P. Donnelly, E. E. Eichler, P. Flicek, S. B. Gabriel, R. A. Gibbs, E. D. Green, M. E. Hurles, B. M. Knoppers, J. O. Korbel, E. S. Lander, C. Lee, H. Lehrach, E. R. Mardis, G. T. Marth, G. A. McVean, D. A. Nickerson, J. P. Schmidt, S. T. Sherry, J. Wang, R. K. Wilson, R. A. Gibbs, E. Boerwinkle, H. Doddapaneni, Y. Han, V. Korchina, C. Kovar, S. Lee, D. Muzny, J. G. Reid, Y. Zhu, J. Wang, Y. Chang, Q. Feng, X. Fang, X. Guo, M. Jian, H. Jiang, X. Jin, T. Lan, G. Li, J. Li, Y. Li, S. Liu, X. Liu, Y. Lu, X. Ma, M. Tang, B. Wang, G. Wang, H. Wu, R. Wu, X. Xu, Y. Yin, D. Zhang, W. Zhang, J. Zhao, M. Zhao, X. Zheng, E. S. Lander, D. M. Altshuler, S. B. Gabriel, N. Gupta, N. Gharani, L. H. Toji, N. P. Gerry, A. M. Resch, P. Flicek, J. Barker, L. Clarke, L. Gil, S. E. Hunt, G. Kelman, E. Kulesha, R. Leinonen, W. M. McLaren, R. Radhakrishnan, A. Roa, D. Smirnov, R. E. Smith, I. Streeter, A. Thormann, I. Toneva, B. Vaughan, X. Zheng-Bradley, D. R. Bentley, R. Grocock, S. Humphray, T. James, Z. Kingsbury, H. Lehrach, R. Sudbrak, M. W. Albrecht, V. S. Amstislavskiy, T. A. Borodina, M. Lienhard, F. Mertes, M. Sultan, B. Timmermann, M.-L. Yaspo, E. R. Mardis, R. K. Wilson, L. Fulton, R. Fulton, S. T. Sherry, V. Ananiev, Z. Belaia, D. Beloslyudtsev, N. Bouk, C. Chen, D. Church, R. Cohen, C. Cook, J. Garner, T. Hefferon, M. Kimelman, C. Liu, J. Lopez, P. Meric, C. O’Sullivan, Y. Ostapchuk, L. Phan, S. Ponomarov, V. Schneider, E. Shekhtman, K. Sirotkin, D. Slotta, H. Zhang, G. A. McVean, R. M. Durbin, S. Balasubramaniam, J. Burton, P. Danecek, T. M. Keane, A. Kolb-Kokocinski, S. McCarthy, J. Stalker, M. Quail, J. P. Schmidt, C. J. Davies, J. Gollub, T. Webster, B. Wong, Y. Zhan, A. Auton, C. L. Campbell, Y. Kong, A. Marcketta, R. A. Gibbs, F. Yu, L. Antunes, M. Bainbridge, D. Muzny, A. Sabo, Z. Huang, J. Wang, L. J. M. Coin, L. Fang, X. Guo, X. Jin, G. Li, Q. Li, Y. Li, Z. Li, H. Lin, B. Liu, R. Luo, H. Shao, Y. Xie, C. Ye, C. Yu, F. Zhang, H. Zheng, H. Zhu, C. Alkan, E. Dal, F. Kahveci, G. T. Marth, E. P. Garrison, D. Kural, W.-P. Lee, W. Fung Leong, M. Stromberg, A. N. Ward, J. Wu, M. Zhang, M. J. Daly, M. A. DePristo, R. E. Handsaker, D. M. Altshuler, E. Banks, G. Bhatia, G. del Angel, S. B. Gabriel, G. Genovese, N. Gupta, H. Li, S. Kashin, E. S. Lander, S. A. McCarroll, J. C. Nemesh, R. E. Poplin, S. C. Yoon, J. Lihm, V. Makarov, A. G. Clark, S. Gottipati, A. Keinan, J. L. Rodriguez-Flores, J. O. Korbel, T. Rausch, M. H. Fritz, A. M. Stütz, P. Flicek, K. Beal, L. Clarke, A. Datta, J. Herrero, W. M. McLaren, G. R. S. Ritchie, R. E. Smith, D. Zerbino, X. Zheng-Bradley, P. C. Sabeti, I. Shlyakhter, S. F. Schaffner, J. Vitti, D. N. Cooper, E. V. Ball, P. D. Stenson, D. R. Bentley, B. Barnes, M. Bauer, R. Keira Cheetham, A. Cox, M. Eberle, S. Humphray, S. Kahn, L. Murray, J. Peden, R. Shaw, E. E. Kenny, M. A. Batzer, M. K. Konkel, J. A. Walker, D. G. MacArthur, M. Lek, R. Sudbrak, V. S. Amstislavskiy, R. Herwig, E. R. Mardis, L. Ding, D. C. Koboldt, D. Larson, K. Ye, S. Gravel, A. Swaroop, E. Chew, T. Lappalainen, Y. Erlich, M. Gymrek, T. Frederick Willems, J. T. Simpson, M. D. Shriver, J. A. Rosenfeld, C. D. Bustamante, S. B. Montgomery, F. M. De La Vega, J. K. Byrnes, A. W. Carroll, M. K. DeGorter, P. Lacroute, B. K. Maples, A. R. Martin, A. Moreno-Estrada, S. S. Shringarpure, F. Zakharia, E. Halperin, Y. Baran, C. Lee, E. Cerveira, J. Hwang, A. Malhotra, D. Plewczynski, K. Radew, M. Romanovitch, C. Zhang, F. C. L. Hyland, D. W. Craig, A. Christoforides, N. Homer, T. Izatt, A. A. Kurdoglu, S. A. Sinari, K. Squire, S. T. Sherry, C. Xiao, J. Sebat, D. Antaki, M. Gujral, A. Noor, K. Ye, E. G. Burchard, R. D. Hernandez, C. R. Gignoux, D. Haussler, S. J. Katzman, W. James Kent, B. Howie, A. Ruiz-Linares, E. T. Dermitzakis, S. E. Devine, G. R. Abecasis, H. Min Kang, J. M. Kidd, T. Blackwell, S. Caron, W. Chen, S. Emery, L. Fritsche, C. Fuchsberger, G. Jun, B. Li, R. Lyons, C. Scheller, C. Sidore, S. Song, E. Sliwerska, D. Taliun, A. Tan, R. Welch, M. Kate Wing, X. Zhan, P. Awadalla, A. Hodgkinson, Y. Li, X. Shi, A. Quitadamo, G. Lunter, G. A. McVean, J. L. Marchini, S. Myers, C. Churchhouse, O. Delaneau, A. Gupta-Hinch, W. Kretzschmar, Z. Iqbal, I. Mathieson, A. Menelaou, A. Rimmer, D. K. Xifara, T. K. Oleksyk, Y. Fu, X. Liu, M. Xiong, L. Jorde, D. Witherspoon, J. Xing, E. E. Eichler, B. L. Browning, S. R. Browning, F. Hormozdiari, P. H. Sudmant, E. Khurana, R. M. Durbin, M. E. Hurles, C. Tyler-Smith, C. A. Albers, Q. Ayub, S. Balasubramaniam, Y. Chen, V. Colonna, P. Danecek, L. Jostins, T. M. Keane, S. McCarthy, K. Walter, Y. Xue, M. B. Gerstein, A. Abyzov, S. Balasubramanian, J. Chen, D. Clarke, Y. Fu, A. O. Harmanci, M. Jin, D. Lee, J. Liu, X. Jasmine Mu, J. Zhang, Y. Zhang, Y. Li, R. Luo, H. Zhu, C. Alkan, E. Dal, F. Kahveci, G. T. Marth, E. P. Garrison, D. Kural, W.-P. Lee, A. N. Ward, J. Wu, M. Zhang, S. A. McCarroll, R. E. Handsaker, D. M. Altshuler, E. Banks, G. del Angel, G. Genovese, C. Hartl, H. Li, S. Kashin, J. C. Nemesh, K. Shakir, S. C. Yoon, J. Lihm, V. Makarov, J. Degenhardt, J. O. Korbel, M. H. Fritz, S. Meiers, B. Raeder, T. Rausch, A. M. Stütz, P. Flicek, F. Paolo Casale, L. Clarke, R. E. Smith, O. Stegle, X. Zheng-Bradley, D. R. Bentley, B. Barnes, R. Keira Cheetham, M. Eberle, S. Humphray, S. Kahn, L. Murray, R. Shaw, E.-W. Lameijer, M. A. Batzer, M. K. Konkel, J. A. Walker, L. Ding, I. Hall, K. Ye, P. Lacroute, C. Lee, E. Cerveira, A. Malhotra, J. Hwang, D. Plewczynski, K. Radew, M. Romanovitch, C. Zhang, D. W. Craig, N. Homer, D. Church, C. Xiao, J. Sebat, D. Antaki, V. Bafna, J. Michaelson, K. Ye, S. E. Devine, E. J. Gardner, G. R. Abecasis, J. M. Kidd, R. E. Mills, G. Dayama, S. Emery, G. Jun, X. Shi, A. Quitadamo, G. Lunter, G. A. McVean, K. Chen, X. Fan, Z. Chong, T. Chen, D. Witherspoon, J. Xing, E. E. Eichler, M. J. Chaisson, F. Hormozdiari, J. Huddleston, M. Malig, B. J. Nelson, P. H. Sudmant, N. F. Parrish, E. Khurana, M. E. Hurles, B. Blackburne, S. J. Lindsay, Z. Ning, K. Walter, Y. Zhang, M. B. Gerstein, A. Abyzov, J. Chen, D. Clarke, H. Lam, X. Jasmine Mu, C. Sisu, J. Zhang, Y. Zhang, R. A. Gibbs, F. Yu, M. Bainbridge, D. Challis, U. S. Evani, C. Kovar, J. Lu, D. Muzny, U. Nagaswamy, J. G. Reid, A. Sabo, J. Yu, X. Guo, W. Li, Y. Li, R. Wu, G. T. Marth, E. P. Garrison, W. Fung Leong, A. N. Ward, G. del Angel, M. A. DePristo, S. B. Gabriel, N. Gupta, C. Hartl, R. E. Poplin, A. G. Clark, J. L. Rodriguez-Flores, P. Flicek, L. Clarke, R. E. Smith, X. Zheng-Bradley, D. G. MacArthur, E. R. Mardis, R. Fulton, D. C. Koboldt, S. Gravel, C. D. Bustamante, D. W. Craig, A. Christoforides, N. Homer, T. Izatt, S. T. Sherry, C. Xiao, E. T. Dermitzakis, G. R. Abecasis, H. Min Kang, G. A. McVean, M. B. Gerstein, S. Balasubramanian, L. Habegger, H. Yu, P. Flicek, L. Clarke, F. Cunningham, I. Dunham, D. Zerbino, X. Zheng-Bradley, K. Lage, J. Berg Jespersen, H. Horn, S. B. Montgomery, M. K. DeGorter, E. Khurana, C. Tyler-Smith, Y. Chen, V. Colonna, Y. Xue, M. B. Gerstein, S. Balasubramanian, Y. Fu, D. Kim, A. Auton, A. Marcketta, R. Desalle, A. Narechania, M. A. Wilson Sayres, E. P. Garrison, R. E. Handsaker, S. Kashin, S. A. McCarroll, J. L. Rodriguez-Flores, P. Flicek, L. Clarke, X. Zheng-Bradley, Y. Erlich, M. Gymrek, T. Frederick Willems, C. D. Bustamante, F. L. Mendez, G. David Poznik, P. A. Underhill, C. Lee, E. Cerveira, A. Malhotra, M. Romanovitch, C. Zhang, G. R. Abecasis, L. Coin, H. Shao, D. Mittelman, C. Tyler-Smith, Q. Ayub, R. Banerjee, M. Cerezo, Y. Chen, T. W. Fitzgerald, S. Louzada, A. Massaia, S. McCarthy, G. R. Ritchie, Y. Xue, F. Yang, R. A. Gibbs, C. Kovar, D. Kalra, W. Hale, D. Muzny, J. G. Reid, J. Wang, X. Dan, X. Guo, G. Li, Y. Li, C. Ye, X. Zheng, D. M. Altshuler, P. Flicek, L. Clarke, X. Zheng-Bradley, D. R. Bentley, A. Cox, S. Humphray, S. Kahn, R. Sudbrak, M. W. Albrecht, M. Lienhard, D. Larson, D. W. Craig, T. Izatt, A. A. Kurdoglu, S. T. Sherry, C. Xiao, D. Haussler, G. R. Abecasis, G. A. McVean, R. M. Durbin, S. Balasubramaniam, T. M. Keane, S. McCarthy, J. Stalker, A. Chakravarti, B. M. Knoppers, G. R. Abecasis, K. C. Barnes, C. Beiswanger, E. G. Burchard, C. D. Bustamante, H. Cai, H. Cao, R. M. Durbin, N. P. Gerry, N. Gharani, R. A. Gibbs, C. R. Gignoux, S. Gravel, B. Henn, D. Jones, L. Jorde, J. S. Kaye, A. Keinan, A. Kent, A. Kerasidou, Y. Li, R. Mathias, G. A. McVean, A. Moreno-Estrada, P. N. Ossorio, M. Parker, A. M. Resch, C. N. Rotimi, C. D. Royal, K. Sandoval, Y. Su, R. Sudbrak, Z. Tian, S. Tishkoff, L. H. Toji, C. Tyler-Smith, M. Via, Y. Wang, H. Yang, L. Yang, J. Zhu, W. Bodmer, G. Bedoya, A. Ruiz-Linares, Z. Cai, Y. Gao, J. Chu, L. Peltonen, A. Garcia-Montero, A. Orfao, J. Dutil, J. C. Martinez-Cruzado, T. K. Oleksyk, K. C. Barnes, R. A. Mathias, A. Hennis, H. Watson, C. McKenzie, F. Qadri, R. LaRocque, P. C. Sabeti, J. Zhu, X. Deng, P. C. Sabeti, D. Asogun, O. Folarin, C. Happi, O. Omoniwa, M. Stremlau, R. Tariyal, M. Jallow, F. Sisay Joof, T. Corrah, K. Rockett, D. Kwiatkowski, J. Kooner, T. Tịnh Hiê`n, S. J. Dunstan, N. Thuy Hang, R. Fonnie, R. Garry, L. Kanneh, L. Moses, P. C. Sabeti, J. Schieffelin, D. S. Grant, C. Gallo, G. Poletti, D. Saleheen, A. Rasheed, L. D. Brooks, A. L. Felsenfeld, J. E. McEwen, Y. Vaydylevich, E. D. Green, A. Duncanson, M. Dunn, J. A. Schloss, J. Wang, H. Yang, A. Auton, L. D. Brooks, R. M. Durbin, E. P. Garrison, H. Min Kang, J. O. Korbel, J. L. Marchini, S. McCarthy, G. A. McVean, G. R. Abecasis, A global reference for human genetic variation. *Nature* **526**, 68–74 (2015).

38.	K. Prüfer, F. Racimo, N. Patterson, F. Jay, S. Sankararaman, S. Sawyer, A. Heinze, G. Renaud, P. H. Sudmant, C. de Filippo, H. Li, S. Mallick, M. Dannemann, Q. Fu, M. Kircher, M. Kuhlwilm, M. Lachmann, M. Meyer, M. Ongyerth, M. Siebauer, C. Theunert, A. Tandon, P. Moorjani, J. Pickrell, J. C. Mullikin, S. H. Vohr, R. E. Green, I. Hellmann, P. L. F. Johnson, H. Blanche, H. Cann, J. O. Kitzman, J. Shendure, E. E. Eichler, E. S. Lein, T. E. Bakken, L. V Golovanova, V. B. Doronichev, M. V Shunkov, A. P. Derevianko, B. Viola, M. Slatkin, D. Reich, J. Kelso, S. Pääbo, The complete genome sequence of a Neanderthal from the Altai Mountains. *Nature* **505**, 43–49 (2014).

55.	K. Morisaki, *Kyusekki Shakai No Jinrui Seitaigaku* (Douseisha publishing, Tokyo, 2022).
